# Novel but stable endosymbionts have contrasting effects on aphid dispersal and plant feeding damage in the cereal pest *Diuraphis noxia*

**DOI:** 10.64898/2026.03.29.715140

**Authors:** Xinyue Gu, Alex Gill, Qiong Yang, Perran A. Ross, Lucy Hayward, Monica Stelmach, Paul A. Umina, Sheik Nadeem Elahee Doomun, Mel Berran, Lauren Coakley, Sonia Sharma, Ary A. Hoffmann

## Abstract

Endosymbiotic bacteria can affect many ecological attributes of their insect hosts, including (in herbivorous insects) how they interact with plants where they feed. This raises the issue of whether deliberate endosymbiont introductions could be used to decrease crop damage caused by insect pests. Here we investigate how transinfecting *Rickettsiella viridis* and *Regiella insecticola* endosymbionts into a novel pest aphid host, the Russian wheat aphid (*Diuraphis noxia*), influences population growth, alate production, dispersal ability and crop damage. Both the *Rickettsiella* (originating from pea aphids) and *Regiella* (from green peach aphids) were stably maintained in their new host where they had contrasting effects. *Rickettsiella* increased the severity of aphid feeding damage on wheat and barley, resulting in greater leaf loss, chlorotic streaking, and higher aphid populations, whereas *Regiella* reduced aphid population growth and the severity of feeding damage by aphids. Their effects on dispersal morphology also differed: *Regiella* had no detectable impact on alate incidence, while *Rickettsiella* consistently suppressed wing formation in small cages, and in larger mesocosms with multiple wheat plants, this endosymbiont suppressed dispersal. Endosymbiont-mediated changes in feeding damage did not involve the main plant immune response pathways: transinfected and wild type aphids induced similar levels of jasmonic acid, jasmonic acid-isoleucine, and salicylic acid in plant tissues, even though these plant defenses were strongly activated during aphid feeding. Novel endosymbionts can therefore modulate the severity of plant feeding damage by aphids as well as influencing aphid dispersal. Potential applications in controlling pest *D. noxia* populations are discussed.

**Significance statement:** Endosymbiotic bacteria that live within insect cells can have wide-ranging effects on the reproduction and fitness of their insect hosts in different environments. In herbivorous insects, this includes effects on host plant use. Here we test if novel endosymbionts in a pest aphid, the Russian wheat aphid, might be used to decrease crop damage and dispersal under controlled laboratory conditions. We show that the damage caused to wheat and barley plants from aphid feeding is modulated by novel but stably transmitted endosymbionts. One endosymbiont (*Rickettsiella*) increased the severity of feeding damage but decreased aphid dispersal, while another (*Regiella*) decreased damage severity without impacting dispersal. These contrasting effects may be associated with changes in aphid population growth and wing formation but were not linked to key plant immune response pathways. We discuss implications of these findings for using endosymbionts in agricultural pest management.

## Introduction

Endosymbionts are heritable bacteria that reside within the cells of their hosts, occurring in more than half of all insect species. They have strong effects on insect phenotypes and can play important roles in shaping insect adaptation and evolutionary trajectories (1–3). Endosymbionts can modulate a range of biological aspects in their host insects including nutrient acquisition (4, 5), thermal tolerance (6–9), reproduction (10–12), resistance to pathogens and natural enemies (13, 14), and plant virus acquisition and transmission (15), thereby influencing the population dynamics of their insect hosts and species interactions.

Beyond influencing insect biology, endosymbionts may also modulate plant defense pathways, including those mediated by salicylic acid (SA) and jasmonic acid (JA), which are central plant responses to herbivory. For instance, in *Bemisia tabaci*, *Rickettsia* endosymbionts transferred into plants activate SA signalling while suppressing JA responses, enhancing herbivore fitness but increasing plant resistance to fungal and viral pathogens (16). This underscores their effects on multitrophic networks involving insects, microbial symbionts, host plants, and other associated pathogens. To date, most studies have focussed on two components, such as endosymbiont–insect or insect–plant interactions, while relatively few have directly examined how all three interact together.

Aphids (Hemiptera: Aphididae) are among the most damaging agricultural pests globally and can be used to investigate this multitrophic network because they host a rich diversity of endosymbionts (17, 18) and share a long co-evolutionary history with some of them dating back approximately 160–280 million years (19). Almost all aphids carry the obligate symbiont *Buchnera aphidicola*, which can provide essential amino acids that aphids cannot obtain from the phloem of plants (5). Some aphids also carry facultative symbionts that exhibit diverse and environmentally dependent effects on hosts (20) and life-history characteristics, such as reproduction (21) and dispersal (22), although impacts on host plant use remain unclear (23). Aphids typically feed on a range of host plant species which include crops that are damaged directly or indirectly through virus transmission (24), leading to substantial economic losses (24–26). Their high reproductive potential, capacity to transmit many economically important plant viruses, and widespread resistance to insecticides further exacerbate their impact (24, 27). Additionally, aphids can reproduce asexually and develop winged progeny, enabling them to relocate to new host plants and disperse widely (28).

Recent studies indicate that endosymbionts may influence aphid–plant–virus interactions. In *Acyrthosiphon pisum*, several endosymbionts may modulate host metabolism and feeding behaviour, thereby influencing virus transmission (29). In *Rhopalosiphum padi*, the transinfected endosymbiont *Regiella insecticola* can interfere with the acquisition and transmission of plant viruses (15). These studies highlight a potential level of complexity of interactions involving endosymbionts beyond direct impacts on host species. For major agricultural pests, it is particularly important to understand how such interactions ultimately shape pest population dynamics and impact crop yield, particularly if there are opportunities to deliberately introduce novel endosymbionts to influence these interactions (30).

Here, we explore endosymbiont-insect-plant interactions following the interspecific transinfection of two endosymbionts, *Rickettsiella viridis* and *Regiella insecticola,* into a global pest of cereal crops, the Russian wheat aphid (*Diuraphis noxia* (Mordwilko)). This follows the successful transinfection of *Rickettsiella* from its natural pea aphid host (*A. pisum*) into the green peach aphid (*Myzus persicae*), where it imposed substantial fitness costs while spreading stably across different host plants through multiple routes (31–33), and the transinfection of *Regiella* from *M. persicae* into its novel aphid host, *R. padi,* with impacts on the transmission of barley yellow dwarf virus (15). While many cross-species aphid transinfections may be unsuccessful or rapidly lost, others can be completely stable (30).

We focused on *D. noxia* due to its importance as an agricultural pest and because native (non-endosymbiotic) bacterial proteins present in *D. noxia* saliva have been linked to aphid feeding damage and virulence to wheat (34). Unlike many other aphid species, *D. noxia* does not transmit plant viruses (35, 36) and rarely carries facultative endosymbionts in the wild (17, 34, 37). *D. noxia* is an invasive pest that has caused major yield losses in wheat and barley, well documented in the USA since its introduction in 1986 (38). Although biotype-specific resistant wheat cultivars can provide temporary control, the continued evolution of *D. noxia* biotypes frequently overcomes host-plant resistance (39, 40). We evaluated the effects of *Rickettsiella* and *Regiella* on aphid fitness, population growth, alate production, feeding damage and plant immune responses (SA and JA levels), as well as endosymbiont spread within populations. By integrating these factors, we investigate how endosymbiont infection influences aphid fitness and aphid–plant interactions, providing insights into the ecological consequences of novel microbial associations and their potential relevance to pest management.

## Results

### *Rickettsiella* and *Regiella* transinfections

We successfully transinfected *Rickettsiella* from the donor *A. pisum* to *D. noxia* via hemolymph microinjection. Of the fifty aphids injected, twelve survived and six of these produced nymphs that tested positive for *Rickettsiella*; one of these lines transmitted *Rickettsiella* to its offspring. Although the *A. pisum* individuals carried both *Serratia symbiotica* and *Rickettsiella*, *Serratia* was not detected in *D. noxia* at G0, whereas *Rickettsiella* was stably maintained at a frequency of 100% for over eighty generations following selection. Similarly, *Regiella* was successfully transinfected from the donor *M. persicae* to *D. noxia*. At G0, eleven of thirty injected aphids tested positive, and three lines exhibited stable transmission to their offspring. These lines were combined at G4, after which *Regiella* persisted at a frequency of 100% for over fifty generations.

### Strain–dependent differences in feeding damage to wheat and barley

*Rickettsiella* and *Regiella* had significant effects on the severity of feeding damage by their host aphids on wheat and barley. On wheat, the *Rickettsiella* strain produced the greatest level of damage, wild type aphids had intermediate effects, and the *Regiella* strain produced the least plant damage (Figure 1A). A repeated-measures ANOVA from Week 2 to Week 3.5 showed a significant main effect of strain on almost all the plant traits, including tiller number (F_2,27_ = 6.280, *p* = 0.006, Figure 1B), leaf number (F_2,27_ = 3.362, *p* = 0.050, Figure 1C), leaf area (F_2,27_ = 7.794, *p* = 0.002, Figure 1D), and overall plant damage (F_2,27_ = 9.099, *p* < 0.001, Figure 1G), with a weaker and non-significant effect on chlorotic streaking (F_2,27_ = 3.181, *p* = 0.067, Figure 1E). All traits exhibited strong effects of time (all *p* < 0.001), reflecting rapid progression of feeding damage across the 3.5 week assessment period. Significant strain × time interactions were detected for aphid infestation level (F_4,54_ = 3.476, p = 0.007, Figure 1F) and leaf area (F_4,54_ = 3.476, p = 0.015, Figure 1D), indicating that the effects of aphid strains changed differently across sampling weeks.

**Figure 1.**
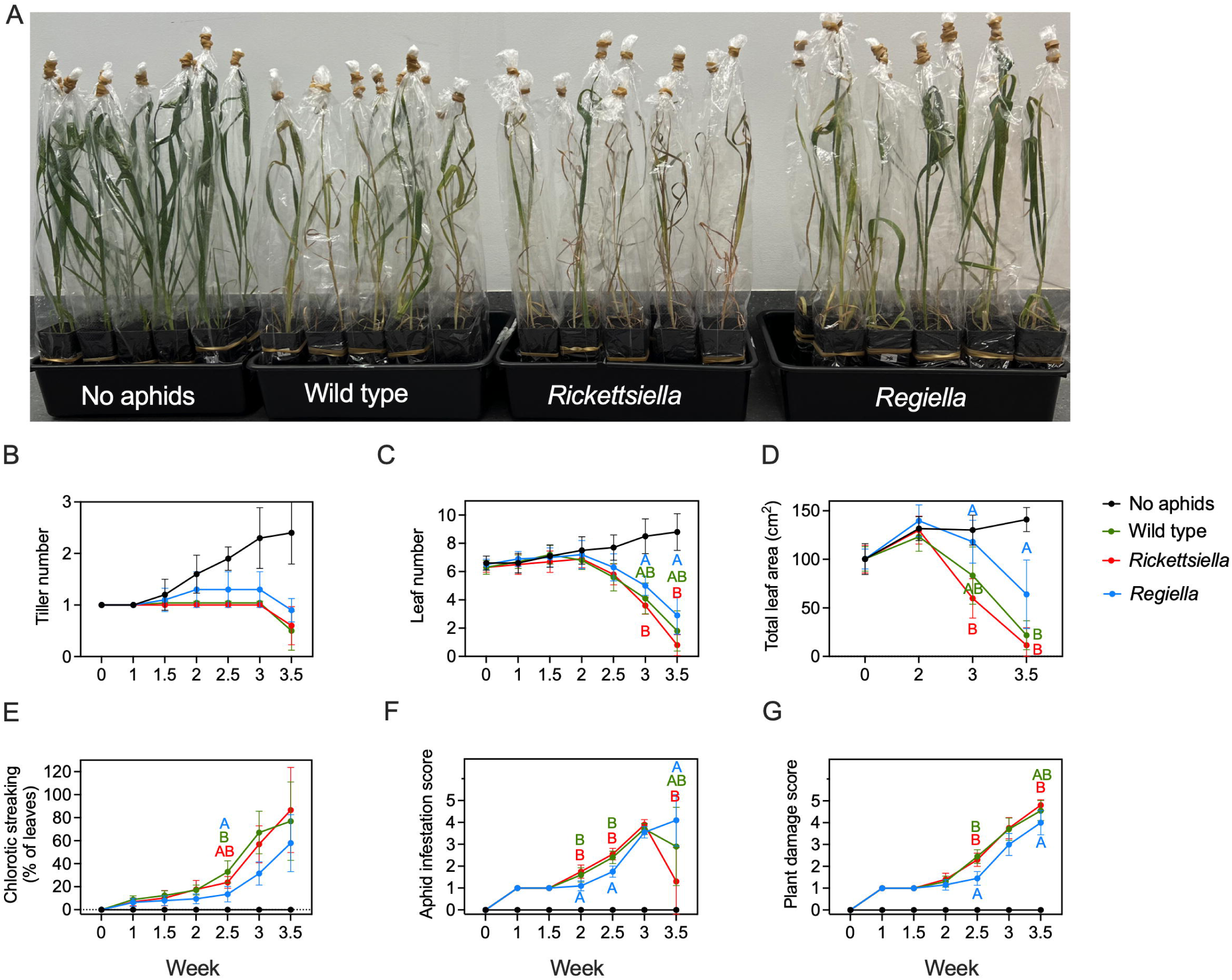
Aphid feeding damage to wheat plants when infested with *Rickettsiella, Regiella* or wild type *D. noxia*. Five 10-day-old apterous females were placed on individual wheat plants. Wheat plants without aphids were included for comparison. (A) Aphid feeding damage to wheat plants when infested with *Rickettsiella*, *Regiella*, wild type *D. noxia* or when lacking aphids at Week 3.5. Each treatment had 10 replicate plants and measured traits were (B) tiller number, (C) leaf number, (D) total leaf area, (E) chlorotic streaking, as well as an (F) aphid infestation score and (G) overall plant damage score. Different letters represent significant pairwise differences between treatments by Bonferroni-adjusted tests. Dots represent means across all replicate wheat plants, and error bars indicate 95% confidence intervals.

Bonferroni–adjusted pairwise comparisons showed that the *Rickettsiella* strain caused more severe plant damage and supported larger aphid populations than the *Regiella* aphid strain on wheat. Specifically, plants infested with *Rickettsiella* aphids had fewer leaves (Week 3: *p* = 0.032; Week 3.5: *p* = 0.028, Figure 1C), reduced leaf area (Week 2.5: *p* = 0.006; Week 3: *p* = 0.002, Figure 1D), higher aphid numbers (Week 2: *p* = 0.004; Week 2.5: *p* < 0.001; Week 3.5: *p* = 0.020, Figure 1F), and greater overall plant damage (Week 2.5: *p* < 0.001; Week 3.5: *p* = 0.022, Figure 1G) compared with plants infested with *Regiella* aphids. Plants infested with the wild type strain also differed from those infested with *Regiella* aphids, with reduced leaf area (Week 3.5: *p* = 0.031, Figure 1D), greater chlorotic streaking (Week 2.5: *p* = 0.031, Figure 1E), higher aphid numbers (Week 2: *p* = 0.031; Week 2.5: *p* = 0.002, Figure 1F), and greater overall plant damage (Week 2.5: *p* < 0.001, Figure 1G).

On barley, feeding damage patterns were similar to those on wheat, although fewer assessments were significantly different between aphid strains. A repeated-measures ANOVA from Week 2 to Week 3 showed a significant main effect of strain on chlorotic streaking (F_2,24_ = 5.016, *p* = 0.015, Figure 1-figure supplement 1E) and overall plant damage (F_2,24_ = 4.142, *p* = 0.028, Figure 1-figure supplement 1G). As on wheat, the *Rickettsiella* strain produced the greatest level of damage, wild type aphids had intermediate effects, and the *Regiella* strain produced the least damage to barley (Figure 1-figure supplement 1A). All plant traits showed a strong effect of time (all *p* < 0.001), reflecting an increase in feeding damage across the 3 week assessment period. However, strain *×* time interactions were not significant (all *p* > 0.095), indicating similar trends in damage over time across strains. Bonferroni–adjusted pairwise comparisons showed greater chlorotic streaking (Week 2: *p* = 0.019, Figure 1-figure supplement 1E) and overall plant damage (Week 2: *p* = 0.034, Figure 1-figure supplement 1G) in plants infested with *Rickettsiella* aphids than with *Regiella* aphids. At Week 3, *Regiella* aphids again tended to produce lower chlorotic streaking (*p* = 0.055) and overall plant damage (*p* = 0.056), although these differences were not statistically significant between strains.

In the barley experiment, we also measured the developmental stages of each aphid strain at Week 3. The *Rickettsiella* aphids had a lower alate frequency than the wild type and *Regiella* aphids (one-way ANOVA: *p* < 0.001, Figure 1-figure supplement 1I), although total adult numbers and total nymph numbers did not differ among strains (all *p* > 0.10, Figure 1-figure supplement 1H & J).

### No effects of Rickettsiella and Regiella on plant defense responses

Using all GC-MS components, PCA’s clearly distinguished metabolic profiles of aphid-feeding areas from non-feeding areas (Figure 2B). However, within either area, there was no clear separation among aphid strains (Figures 2C & 2D), suggesting broadly similar metabolomic profiles among strains of the same aphid clone carrying different symbionts.

**Figure 2.**
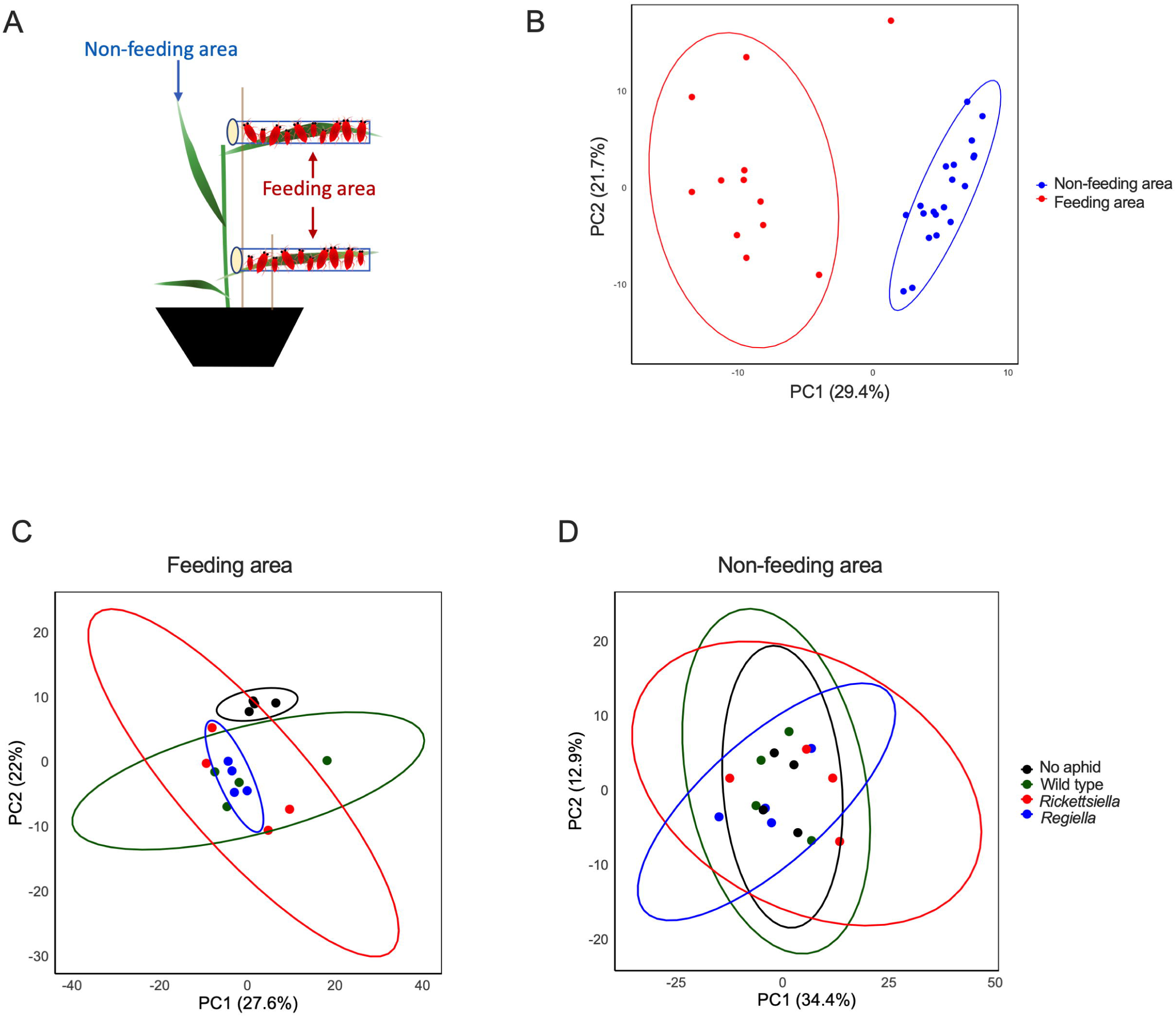
Principal Component Analysis of all GC-MS components showing metabolic profiles: (A) Schematic showing the feeding and non-feeding areas the wheat plants used in this experiment. Sixty age-mixed aphids were divided into two plastic vials. The top and bottom leaves of each plant were inserted into a vial and a sponge plug was placed into the vial openings and left for 7 days. Control plants were treated in same manner, except no aphids were added to the vials. After a 7-day infestation period, all aphids were removed and the plants remained for an additional 7 days before phytohormone analysis was undertaken. (B) Profiles of aphid feeding areas versus non-feeding areas, (C) profiles of the three aphid strains and no-aphid control on the feeding areas, and (D) profiles of the three aphid strains and no-aphid control on the non-feeding areas. Dots show data from individual replicate plants (*n* = 4 per aphid strain), and ovals show 95% confidence regions. For (B), data from areas with no aphids were included as non-feeding areas. For (C) and (D), the same feeding areas with no aphids were included as controls for comparison with the different aphid feeding treatments.

Levels of JA, jasmonic acid-isoleucine (JA-Ile), and SA did not differ significantly among treatments in feeding areas (Figure 2-figure supplement 1A-C) and non-feeding areas (Figure 2-figure supplement 1D-F), indicating that Rickettsiella and Regiella did not induce notable differences in defense responses through the JA and SA pathways.

Heatmap analysis showed largely consistent responses among aphid strains in both feeding areas (Figure 2-figure supplement 2A) and non-feeding areas (Figure 2-figure supplement 2B), with the exception of four metabolites (Figure 2-figure supplement 3). In feeding areas, valine (Figure 2-figure supplement 3A) increased by an average of 1.4-fold, while 3-phenyllactic acid (Figure 2-figure supplement 3B) decreased by an average of 3.3-fold in plants exposed to *Rickettsiella* aphids compared with plants exposed to wild type aphids, suggesting possible alterations in amino acid metabolism and reduction in antimicrobial or defense-related metabolites. In non-feeding areas, urea (Figure 2-figure supplement 3C) decreased by an average of 5.3-fold, while pantothenic acid (Figure 2-figure supplement 3D) increased by an average of 1.5-fold in plants exposed to *Regiella* aphids compared to plants exposed to wild type aphids, potentially indicating altered nitrogen metabolism and enhanced energy and secondary metabolite synthesis caused by *Regiella* aphid feeding.

### No effects of Rickettsiella and Regiella on aphid life history traits

We measured the effects of *Rickettsiella* and *Regiella* on life history traits of their host aphids at two temperatures (19°C and 25°C), as well as body color and body shape/size. For *Rickettsiella,* development time differed between strains (Kruskal-Wallis Test: H_1,62_ =4.405, *p* = 0.036, Figure 3-figure supplement 1A), with *Rickettsiella* aphids developing faster than wild type aphids at 25°C. However, lifetime fecundity (GLM: F_1,118_ = 1.321, *p* = 0.253, Figure 3-figure supplement 1B) and longevity (Cox regression: χ^2^ = 0.153, d.f. = 1, *p* = 0.696, Figure 3-figure supplement 1C) did not differ between *Rickettsiella* and wild type aphids at either temperature, and there was no significant effect on development time at 19°C. Temperature significantly affected development time (Kruskal-Wallis test: H_1,122_ = 86.061, *p* < 0.001) and longevity (Cox regression: χ^2^ = 10.635, d.f. = 1, *p* = 0.001) but not lifetime fecundity (GLM: F_1,118_ = 0.033, *p* = 0.857). Unlike in other aphid species (15, 31, 41), *Rickettsiella* had no effect on body color in *D. noxia* (MANOVA across all color components: *p* = 0.618; t-test: Hue: *p* = 0.795, Figure 3-figure supplement 1D; Saturation: *p* = 0.409, Figure 3-figure supplement 1E; Lightness: *p* = 0.603, Figure 3-figure supplement 1F). Body shape of apterous adults (expressed as the ratio of body length/body width) was also not influenced by *Rickettsiella* infection (t-test: *p* = 0.115, Figure 3-figure supplement 1G).

**Figure 3.**
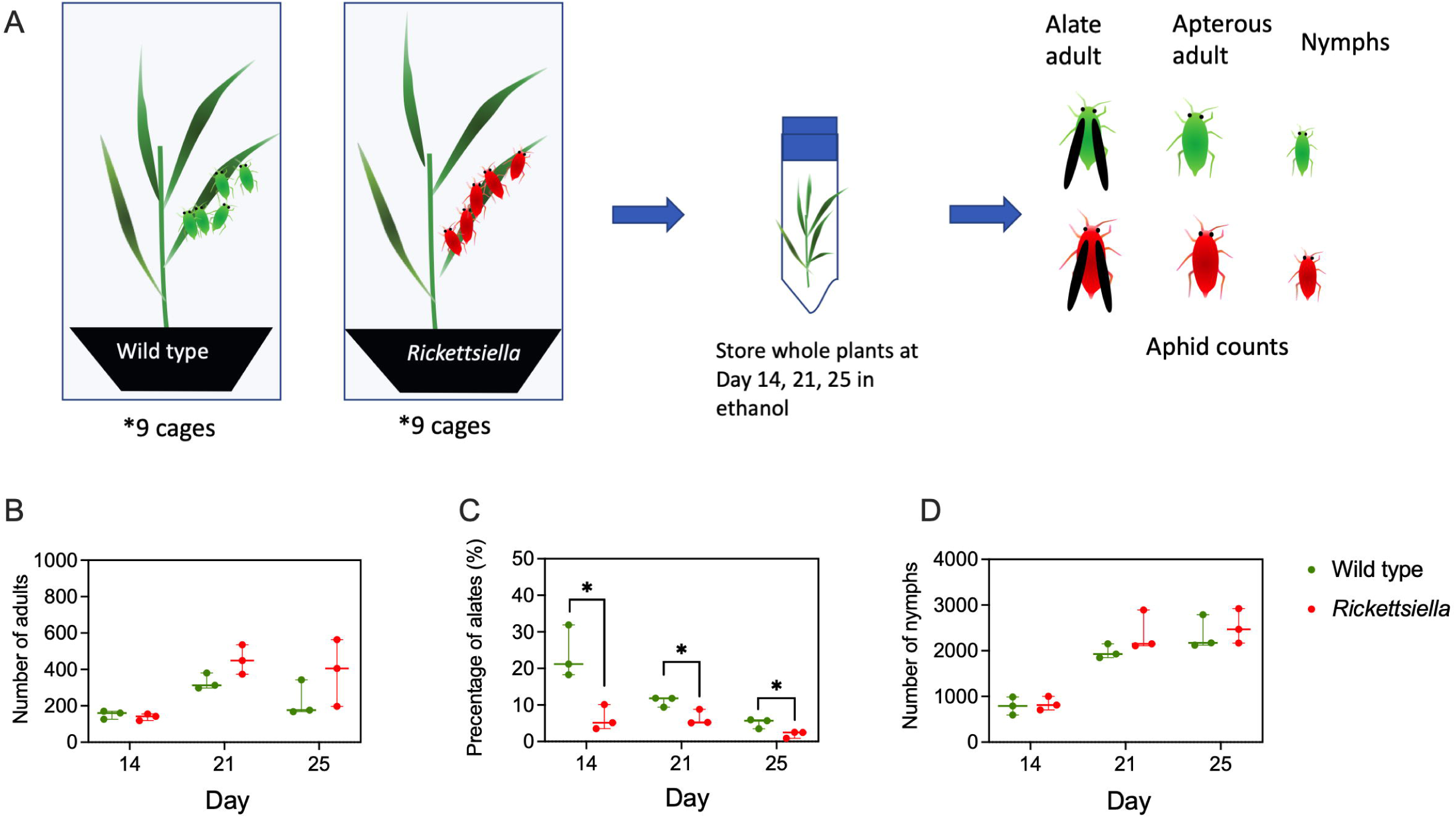
*Rickettsiella* effects on aphid population growth and alate production on whole plants. (A) Experimental design for *Rickettsiella*. Five 14-day-old *Rickettsiella* or wild type apterous female aphids were placed onto wheat plants (GS14). At Day 14, 21, and 25, aphids from three replicate plants per treatment were removed and the number of alate adults, apterous adults and nymphs was recorded. (B) The number of *Rickettsiella* adults (alate plus apterous adults), (C) the percentage of *Rickettsiella* alates and (D) the number of *Rickettsiella* nymphs were used to characterize aphid population growth and alate production over time. An independent sample t-test was used to compare differences between endosymbiont and wild type treatments at each time point, with “*” indicating significant differences (*p* < 0.05).

At 19°C, we measured *Buchnera* and *Rickettsiella* densities, and found no effect of *Rickettsiella* on *Buchnera* density (t-test: *p* = 0.914, Figure 3-figure supplement 1H), suggesting the addition of a novel *Rickettsiella* endosymbiont did not perturb the obligate endosymbiont *Buchnera*. All *Rickettsiella* aphids we screened were positive for *Rickettsiella*, with a median density equivalent to the host control gene, and all wild type aphids screened were negative (Figure 3-figure supplement 1I).

For *Regiella,* there were no effects on development time (Kruskal-Wallis Test: H_1,148_ = 0.449, *p* = 0.503, Figure 4-figure supplement 1A), lifetime fecundity (GLM: F_1,113_ = 5.266, *p* = 0.919, Figure 4-figure supplement 1B), longevity (Cox regression: χ^2^ = 1.719, d.f. = 1, *p* = 0.190, Figure 4-figure supplement 1C), or the body length of apterous adults (F_1,120_ = 2.063, *p* = 0.154, Figure 4-figure supplement 1G) relative to wild type aphids at both 19°C and 25°C. *Regiella* affected body color components except for hue (GLM: Hue: F_1,119_ = 0.658, *p* = 0.544, Figure 4-figure supplement 1D; Saturation: F_1,119_ = 941.134, *p* < 0.001, Figure 4-figure supplement 1E; Lightness: F_1,119_ = 29.579, *p* < 0.001, Figure 4-figure supplement 1F), with *Regiella* aphids being lighter in color and less saturated than wild type aphids. Temperature significantly affected development time (Kruskal-Wallis test: H_1,148_ =103.232, *p* < 0.001), longevity (Cox regression: χ^2^ = 6.904, d.f. = 1, *p* = 0.009), and lifetime fecundity (GLM: F_1,113_ = 12.494, *p* < 0.001), but did not affect body color traits except hue (F_1,119_ = 19.364, *p* < 0.001). No significant strain × temperature interactions were detected for any measured trait (all p > 0.085).

**Figure 4.**
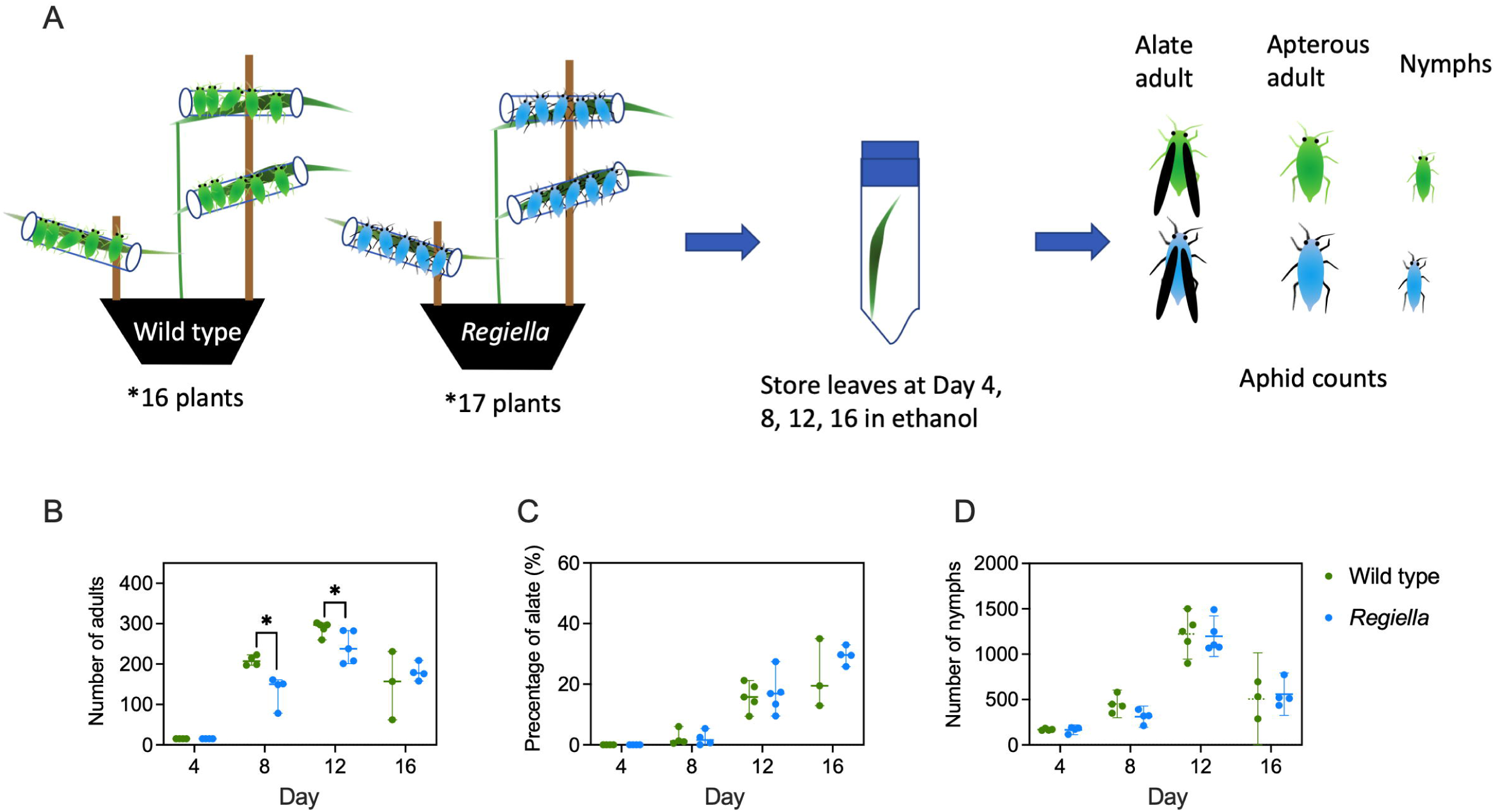
*Regiella* effects on aphid population growth and alate production on whole plants. (A) Experimental design for *Regiella*. Five 14-day-old *Regiella* or wild type apterous female aphids were placed onto each of the top, middle and bottom leaves of wheat plants (GS14). A plastic vial was placed over each leaf and cotton wool inserted into the vial opening to prevent aphid movement. At Days 4, 8, 12, and 16, aphids were removed from replicate plants in each treatment, and the numbers of alate adults, apterous adults, and nymphs were recorded. Four replicate plants per treatment were sampled at Days 4 and 8, five plants infested with *Regiella* and wild type aphids were sampled at Day 12, and the remaining plants were sampled at Day 16. (B) The number of *Regiella* adults (alate plus apterous adults), (C) the percentage of *Regiella* alates and (D) the number of *Regiella* nymphs were used to characterize aphid population growth and alate production over time. The independent sample t-test was used to compare differences between endosymbiont and wild type treatments at each time point, with “*” indicating significant differences (*p* < 0.05). Development stages of aphids on different wheat leaves are shown in Figure 4-figure supplement 3.

At 19°C, *Regiella* did not significantly affect *Buchnera* density (t-test, *p* = 0.075; Figure 4-figure supplement 1H), suggesting the addition of a novel *Regiella* endosymbiont did not perturb the obligate endosymbiont *Buchnera.* All *Regiella* aphids we screened were positive for *Regiella*, with a median density equivalent to the host control gene, and all wild type aphids screened were negative (Figure 4-figure supplement 1I).

It is worth noting the above assays were conducted in cups containing a single wheat plant, whereby aphids were maintained under favourable conditions, with limited environmental stress and without substantial crowding.

### Frequency of Rickettsiella and Regiella in mixed infection cages linked to temperature

We established cages at both 19°C and 25°C containing mixed populations of wild type aphids and aphid strains carrying either *Rickettsiella* or *Regiella,* to investigate population dynamics over time. The proportion of aphids carrying *Rickettsiella* or *Regiella* increased rapidly. *Rickettsiella* reached >75% frequency in most replicate cages by Week 3 at 19°C (Figure 3-figure supplement 2B) and by Week 2 at 25°C (Figure 3-figure supplement 2C). *Regiella* reached 100% frequency at both temperatures by Week 12 (Figure 3-figure supplement 2B & C). At 25°C, *Rickettsiella* maintained a high prevalence through Week 12 (Figure 3-figure supplement 2C), whereas at 19°C its prevalence declined between Weeks 9 and 12 (Figure 3-figure supplement 2B). By contrast, *Regiella* persisted at a higher frequency at 19°C for 30 weeks (Figure 4-figure supplement 2B) but at 25°C its frequency varied among replicate cages between Weeks 18 and 24 (Figure 4-figure supplement 2C).

Horizontal transmission is expected to initially generate aphids with low endosymbiont densities, which may then increase and be vertically transmitted to subsequent generations (31). For *Rickettsiella,* aphids with low endosymbiont densities were common (Figure 3-figure supplement 2F & G) but occurred at relatively lower frequencies at 25°C (Figure 3-figure supplement 2G). High-density infections generally remained stable over time for *Rickettsiella* at 25°C (Figure 3-figure supplement 2E) and for *Regiella* at 19°C (Figure 4-figure supplement 2D), whereas high *Regiella* densities declined at 25°C in some replicate cages from Week 12 (Figure 4-figure supplement 2E & G). The persistence of high-density infections is consistent with minor fitness effects associated with both infections, although any minor fitness disadvantage in infected individuals could also be offset by conversion from low-density infections.

### Different effects of Rickettsiella and Regiella on aphid population growth and alate production on whole plants

We investigated the effects of *Rickettsiella* and *Regiella* on aphid population growth and alate production on whole wheat plants. Aphid populations were established from a small number of aphids and subsequently experienced increasing crowding and density-dependent stress (Figures 3 & 4). Under these conditions, the effects on aphid host fitness were different from those observed in the individual assays. In the *Rickettsiella* experiment, plant numbers were limited (three plants per time point), so analyses focused on Days 21 and 25 where any population growth effects were expected to have built up. There was no effect of strain on total adult numbers (alate + apterous adults) at these two time-points (F_1,8_ = 2.159, *p* = 0.180; Figure 3B), and there was no time *×* strain interaction (F_1,8_ = 0.468, *p* = 0.513). However, *Rickettsiella* significantly reduced alate frequency from Day 14 to Day 25 (GLM: F_1,12_ = 26.434, *p* < 0.001, Figure 3C), and the strain *×* time interaction was also significant (GLM: F_2,12_ = 7.830, *p* = 0.007). The frequency of alates decreased as plant damage increased (Figure 3C) but remained significantly lower for the *Rickettsiella* strain at all sampling points. Total nymph numbers were also higher on plants infested with *Rickettsiella* aphids from Day 21 to Day 25 than on plants infested with wild type aphids (GLM: F_1,8_ = 13.646, *p* = 0.006, Figure 3D), with no strain *×* time interaction (F_1,8_ = 1.070, *p* = 0.331).

In the *Regiella* experiment, we used a slightly different design whereby aphids were introduced at multiple positions on individual wheat plants (see “*Population growth and alate production on whole plants*“). Here, we focused on the period after 4 days when aphid numbers were still building up. Adult *Regiella* aphids were significantly fewer than wild type adults on Days 8 and 12 (F_1,14_ = 14.049, *p* = 0.002, Figure 4B), but this difference was not evident at Day 16 (F_1,19_ = 1.045, *p* = 0.320). The time *×* strain interaction for adult numbers from Day 8 to Day 16 was significant (F_2,19_ = 3.868, *p* = 0.039, Figure 4B). Aphid strain did not affect alate frequency (Day 8 to 12: F_1,14_ =0.040, *p* = 0.844; Day 16: F_1,19_ =1.385, *p* = 0.254; Figure 4C) or the total nymph number (Day 8 to 12: F_1,14_ = 4.315, *p* = 0.057; Day 16: F_1,19_ = 0.708, *p* = 0.411; Figure 4D). Time *×* strain interactions were also not significant for alate frequency (F_2,19_ = 0.893, *p* = 0.426, Figure 4C) or total nymph number (F_2,19_ = 0.599, *p* = 0.622, Figure 4D) from Day 8 to Day 16.

### Rickettsiella reduces alate production and increases feeding damage to wheat plants in mesocosms

We further tested the effects of *Rickettsiella* on aphid dispersal ability and feeding damage to wheat plants in a mesocosm experiment in which *Rickettsiella* and wild type aphids were released into the same enclosures. Patterns of feeding damage were consistent with our earlier experiments using individual cages (Figure 1), with greater feeding damage over time in the *Rickettsiella* treatment than in the wild type treatment (Figure 5). The clearest differences occurred at Week 3.5 on the aphid-release plants, where plants infested with *Rickettsiella* aphids had fewer leaves (t-test: *p* < 0.001, Figure 5B), reduced total leaf area (*p* = 0.050, Figure 5C), higher aphid infestation scores (*p* < 0.001, Figure 5F), and greater overall plant damage (*p* = 0.007, Figure 5G) than plants infested with wild type aphids. Chlorotic streaking also appeared earlier and was greater in plants infested with *Rickettsiella* aphids, with a significant difference at Week 3 (*p* = 0.046, Figure 5D). Similar results were observed for the repeated measures GLM; wheat plants infested with *Rickettsiella* aphids had fewer leaves (GLM: F_1,38_ = 11.368, *p* = 0.002), reduced total leaf area (F_1,38_ = 14.849, *p* < 0.001), greater chlorotic streaking before damage levels converged (Weeks 2-3.5: F_1,31_ = 7.409, *p* = 0.011) and greater overall plant damage (F_1,38_ = 12.414, *p* = 0.001). When a combination of all plant traits was considered in the MANOVAs, feeding damage to wheat plants differed significantly between aphid strains at Week 2 (*p* = 0.023) but differences were marginally non-significant at Week 3 (*p* = 0.050) and Week 4 (*p* = 0.070).

**Figure 5.**
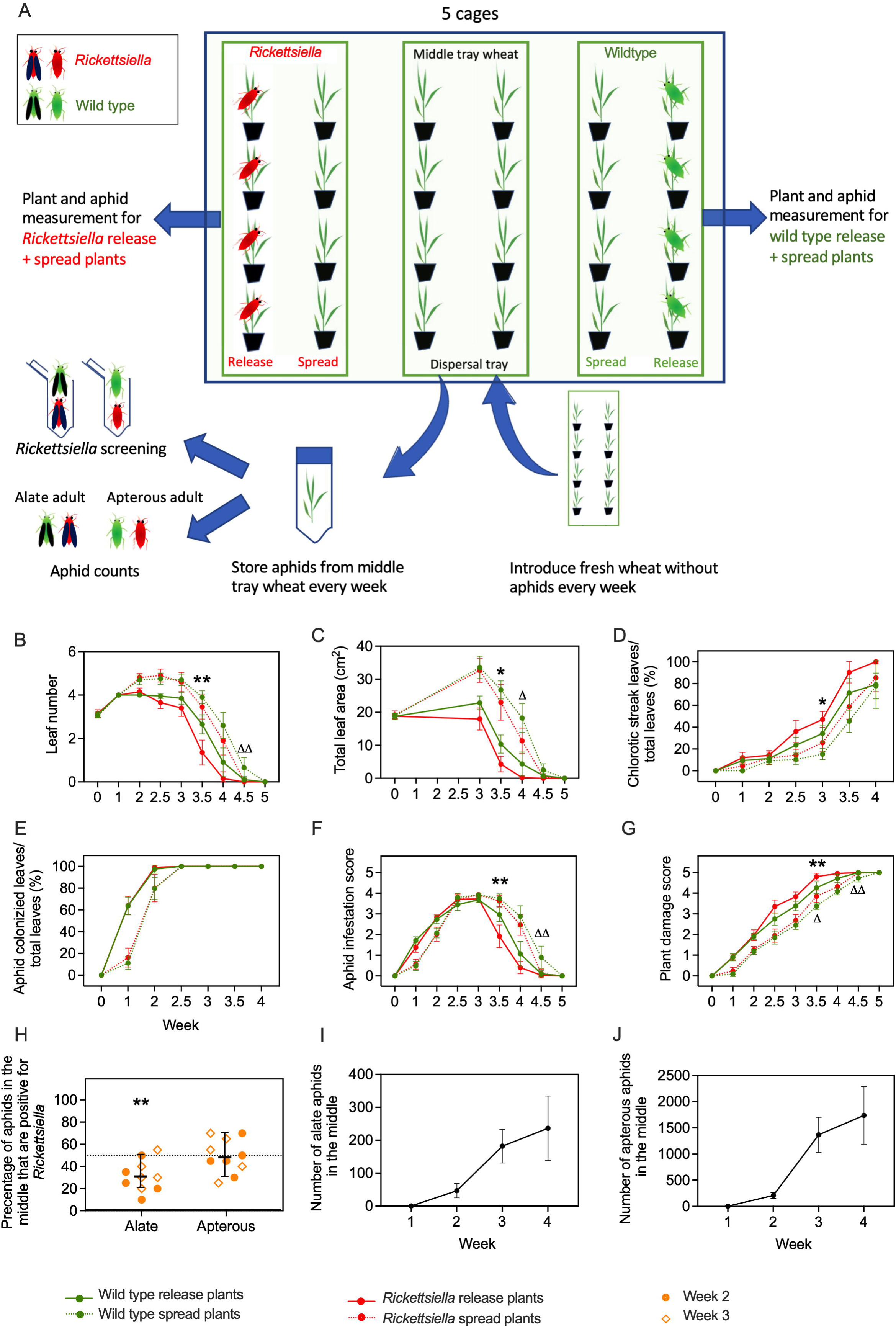
*Rickettsiella* effects on aphid dispersal and plant feeding damage to wheat plants in mesocosms. (A) Experimental design. Populations were initiated with 5 *Rickettsiella* or wild type aphids per plants on four wheat plants (‘aphid-release wheat’) while adjacent plants (‘aphid-spread wheat’) were monitored. All plants were measured for (B) leaf number, (C) total leaf area, (D) chlorotic streaking, (E) aphid colonization rate, as well as (F) an aphid infestation score and (G) overall plant damage score. “*” and “**” represent significant differences at *p* < 0.05 and *p* < 0.01, respectively when comparing *Rickettsiella* and wild type aphids on aphid-release wheat. “*Δ*” and “*ΔΔ*” represent significant differences at *p* < 0.05 and *p* < 0.01, respectively when comparing *Rickettsiella* and wild type aphids on aphid-spread wheat. Aphids collected from the middle tray containing eight wheat plants (“dispersal”) were counted and screened to determine (H) the percentage of alate and apterous adults in the middle that were positive for *Rickettsiella,* (I) the number of alate adults, and (J) the number of apterous adults. Symbols in (H) represent the percentage of 20 alate and 20 apterous adults positive for *Rickettsiella* at Weeks 2 and 3. “**” represents a significant difference at *p* < 0.01 as determined from a paired t-test.

Increased feeding damage from *Rickettsiella* aphids was also evident in the aphid-spread plants, and this effect was most pronounced at Week 4.5. At this time point, there were fewer leaves (t-test: *p* = 0.003, Figure 5B), and greater overall plant damage (*p* = 0.001, Figure 5G) in the *Rickettsiella* treatment compared with the wild type treatment. The *Rickettsiella* treatment also had a higher aphid infestation level (*p* = 0.001, Figure 5F). Total leaf area was also significantly lower in the *Rickettsiella* treatment than in the wild type treatment at Week 4 (*p* = 0.020, Figure 5C). A similar pattern was observed for the repeated measures GLM of the aphid-spread plants; plants infested with *Rickettsiella* aphids had reduced leaf area (F_1,38_ = 4.432, *p* = 0.042) and greater overall plant damage (F_1,38_ = 6.009, *p* = 0.019).

If *Rickettsiella* and wild type aphids exhibited similar population growth and dispersed equally, we would expect a *Rickettsiella* frequency of 50% among aphids sampled from the middle plants. However, at both Weeks 2 and 3, the proportion of alates in the middle tray that tested positive for *Rickettsiella* was significantly below 50% (paired t-test: *p* = 0.002, Figure 5H). In contrast, no significant deviation from 50% was observed for apterous aphids collected from the middle tray (*p* = 0.922). As expected, the numbers of alate (Figure 5I) and apterous adults (Figure 5J) in the middle trays increased over time.

## Discussion

Despite the wide-ranging effects of facultative endosymbionts on aphid phenotypes, relatively few studies have considered endosymbiont–aphid–plant interactions, although enhanced plant immune responses have been reported in some systems (16, 42). Here, we show that transinfections of *Rickettsiella* and *Regiella* in *D. noxia* produced opposing effects on plant damage. *Rickettsiella* increased aphid feeding damage, whereas *Regiella* decreased it in both wheat and barley (Figure 1 & Figure 1-figure supplement 1, Table 1). These effects were evident across multiple plant traits, and were particularly clear for chlorotic streaking, which is an important effect reflecting salivary toxins secreted during *D. noxia* feeding (34).

**Table 1.**
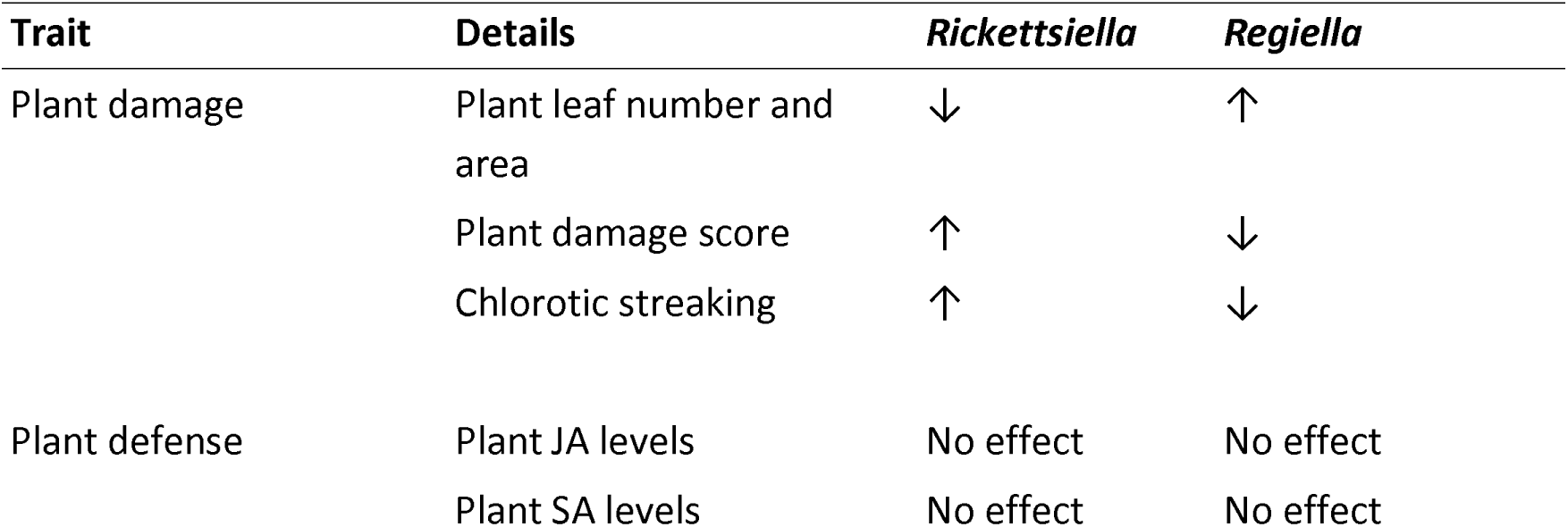
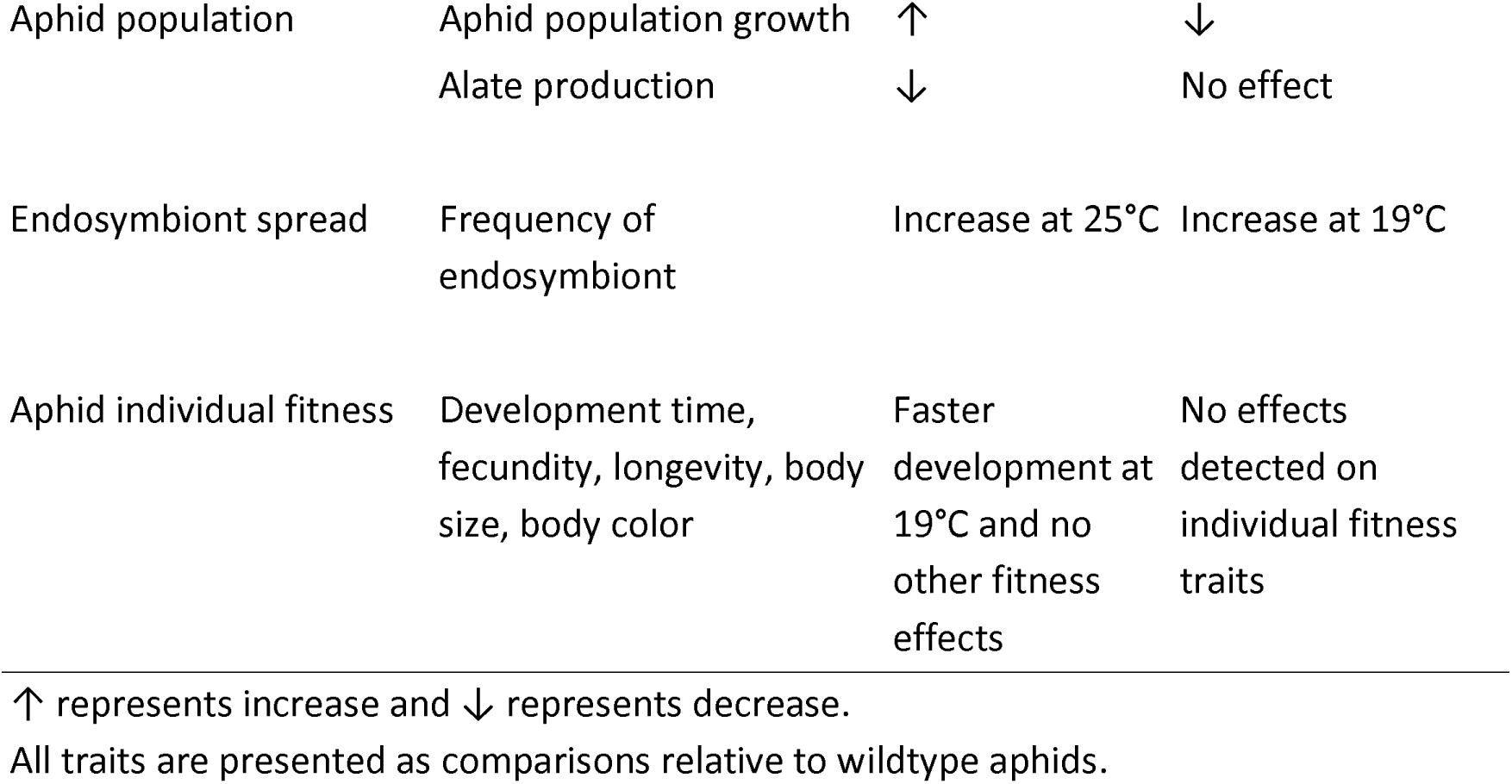
Summary of effects of *Rickettsiella* and *Regiella* transinfections in *Diuraphis noxia*.

However, we did not detect any differences in plant immune responses through the JA or SA pathways, suggesting that the endosymbionts did not reprogram canonical defense signalling in wheat under our experimental conditions.

Despite this, our metabolomic analyses suggest that endosymbionts in *D. noxia* may influence some other aspects of wheat metabolism in a spatially structured way. Within aphid-feeding areas, wheat exposed to *Rickettsiella* aphids showed increased valine and decreased 3-phenyllactic acid. Changes in valine have been reported in plants responding to herbivory and other forms of stress (43–45), while 3-phenyllactic acid has been associated with antimicrobial and defense-related activity (46). In non-feeding areas, wheat exposed to *Regiella* aphids showed reduced urea and increased pantothenic acid. These changes may reflect differences in nitrogen metabolism and allocation (47), and in metabolic processes involving pantothenic acid (48) respectively. However, the present metabolomic data do not establish the functional consequences or causal mechanisms underlying these changes. They indicate that aphids carrying different endosymbionts are associated with some distinct metabolic responses in wheat, including responses that differ between aphid-feeding and non-feeding areas. Our findings are consistent with previous work showing that phloem-feeding insects can induce changes in plant metabolites in response to herbivory (49, 50) and provide a basis for future studies to determine how endosymbionts influence aphid-induced plant responses and contribute to contrasting plant phenotypes.

Apart from inducing metabolic changes, endosymbiont effects on plant damage may reflect population-level effects rather than (or in addition to) direct plant-endosymbiont interactions. In our single plant experiments, where aphids were maintained under relatively favourable conditions, with limited environmental stress and without substantial crowding, we detected few fitness effects attributable to endosymbionts. In contrast, our population growth experiments, in which populations experienced increasing crowding and density-dependent stress, found that *D. noxia* carrying *Rickettsiella* had a greater population size, whereas those carrying *Regiella* showed a reduction in population size. These results suggest endosymbiont-mediated fitness effects are context-dependent and may become more pronounced as aphid population density increases and density-dependent stress develops. Thus, relatively modest effects observed at the individual level can potentially translate into meaningful differences at the population level, which could contribute to differences in feeding pressure and symptom development. The direction of the *Rickettsiella* effects in *D. noxia* differs from patterns reported in transinfected *M. persicae* (31, 32), underscoring that fitness consequences of facultative symbionts are host specific.

Both endosymbionts persisted and spread rapidly in mixed cages, with dynamics partly influenced by temperature. *Rickettsiella* frequency declined late in the 19°C cages, whereas *Regiella* ultimately reached fixation at both temperatures but showed greater among-replicate variability at 25°C later in the experiment. These outcomes are potentially shaped by both vertical transmission and horizontal transmission through plant tissue (16, 51, 52) and/or transfer via aphid contact or honeydew (53, 54). Similar patterns of context-dependent symbiont effects have been reported in other aphid species (55) and may be relevant to variability in aphid endosymbiont frequencies under field conditions where additional factors such as parasitism interactions can be important (56–58). Although we did not directly document conversion of low endosymbiont titre to high titre offspring as previously demonstrated for *Rickettsiella* in *M. persicae* (31), we observed a substantial increase in the incidence of high-titre infections of both endosymbionts following periods of increase in the total incidence of infections, which suggests that this conversion may be ongoing. However serial bottlenecks during transfers likely generated a high level of variability among replicates.

A noteworthy result is that *Rickettsiella* reduced alate production in *D. noxia*, whereas *Regiella* did not. Wing polymorphism is an important adaptative response that facilitates aphid dispersal in response to crowding, host quality, and other environmental cues (59). Previous studies have shown that *Serratia* can promote wing development in *A. pisum* that enhances its dispersal (22) and that suppression of *Buchnera* through antibiotic treatment can reduce alate production in *Sitobion avenae* (60). We have also found that transinfection of *Rickettsiella* into *R. padi* reduced both host fitness and alate production, further supporting an effect of this endosymbiont in dispersal-related traits across aphid hosts (61). The suppression of wing development and alate dispersal by *Rickettsiella* in *D. noxia* across experiments in wheat and barley was apparent even when total adult numbers were high, indicating that this effect may persist irrespective of crowding. Future work could directly measure the effects of *Rickettsiella* on wing formation as undertaken in other studies (e.g. 22). Given wing formation is highly sensitive to environmental conditions (62), it would also be interesting to complete multiple-generation experiments in large arenas at different temperatures and using plants experiencing different stress levels.

It would be worthwhile undertaking further work to better understand the mechanisms underlying the symbiont-associated differences in plant damage. Aphid feeding behavior could be examined using electrical penetration graph (EPG) assays to test whether symbionts alter probing dynamics, phloem access, or toxic components in aphid saliva. Recent work has shown that endosymbionts can alter probing and feeding behaviour (63, 64) and the composition of salivary proteins (65) in aphids. Virulent biotypes of *D. noxia* are known to produce more additional energy than less virulent biotypes under stressful conditions based on measures of the NAD+/NADH ratio (66), and such differences could also be impacted by endosymbionts (67). Luna *et al.* (2018) identified bacterial proteins in *D. noxia* saliva that facilitates virulence to wheat. Apart from bacteria, recent studies have also showed non-coding RNA (68) and epigenetic effects (69) that alter virulence to host plants. Comparing salivary toxins and regulatory pathways among *Rickettsiella*, wild type and *Regiella* aphids could therefore help identify how symbionts influence damage-related traits and potentially contribute to understanding how *D. noxia* develops new virulent populations despite limited genetic diversity (70). It would also be worth looking at other methods to detect and quantify endosymbiont levels. Although PCR and qPCR are widely used for this purpose, factors such as primer specificity, amplification efficiency, and sequence variation can affect assay accuracy and result in false negatives (71). Complementary approaches, such as flow cytometry (72) and SymbiQuant (73), could be valuable for validating and improving estimates of symbiont abundance in future studies.

The transinfected strains could be useful in aphid management depending on the ecology of the pest populations. If *Regiella* reduces plant damage by suppressing *D. noxia* population growth, it could potentially lessen crop impact once established in populations. Conversely, *Rickettsiella* establishment may limit aphid spread. These potential benefits need to be weighed against risks which will likely depend on the cropping situation. In related work, we have found that greater plant damage from *Rickettsiella* extends beyond wheat and barley to summer ‘green-bridge’ hosts such as barley grass and brome grass (74, 75), which could help reduce *D. noxia* populations between winter-cropping seasons. The combined effects of *Rickettsiella* on dispersal and green bridge hosts could offset any increases in crop damage.

Implementation of endosymbiont-based programs will also depend on clarifying other risks. Current data suggest that *Rickettsiella* transinfections have little impact on pesticide tolerance in *D. noxia* and other aphids (76), an important risk dimension given that chemicals remain the major way *D. noxia* are managed in agricultural fields in Australia (75). Experimental evaluations also need to be extended to a broader range of wheat varieties, particularly resistant lines, although this is not currently feasible locally because only *D. noxia*–susceptible varieties are available in Australia. Additionally, it would be worthwhile exploring effects in other *D. noxia* clones given that symbiont effects can vary with aphid genotype (58, 77) although only a single lineage of *D. noxia* is likely to be present in Australia (Holland et al., unpublished data) which likely reflects the recent incursion of this species from overseas (78). Nevertheless, the current results suggest that transinfections can affect *D. noxia* behaviour and plant interactions in interesting ways that might be useful in pest management.

## Materials and methods

### Aphid strains and maintenance

Aphids were cultured using a cup culturing method {Yu, 2025 #137}. Plant material was inserted into cups with a reservoir of water to support the plant and aphid population. The host plant for *D. noxia* was wheat (*Triticum aestivum* cv. Trojan), for *M. persicae* it was bok choy (*Brassica rapa* subsp. chinensis var. baby bok), and for *A. pisum* it was lucerne (*Medicago sativa*, cv. Sequel). Aphids were moved to new cups with fresh leaf material twice a week and maintained in temperature-controlled cabinets at 19 (± 1)°C with a 16:8h L:D photoperiod. All plant material was grown in insect-proof BugDorm cages (93 x 47.5 x 47.5cm, mesh 160 µm aperture, Australian Entomological Supplies, Australia) within a shade-house supported with plant growth lights (40W Grow Saber LED 6500K, 1200mm length) set to a 16:8h L:D photoperiod. Wheat and bok choy plants were grown for ∼ 6 weeks before use, while lucerne plants were ∼ 4.5 weeks old before use. For our experiments, wheat plants (∼ 5 weeks old) and barley plants (*Hordeum vulgare* var. La Trobe ∼ 5 weeks old) were grown under the same conditions.

For donor species that carried the facultative endosymbionts, a single clone of *A. pisum* was collected from Tintinara, South Australia, Australia (GPS: -35.95S, 140.14E), which carried both *Serratia symbiotica* and *Rickettsiella viridis*, and a single *M. persicae* clone was collected from Burnley, Victoria, Australia (GPS: -35.83S, 145.02E), which carried *Regiella insecticola*. For the recipient *D. noxia,* we used a single clone collected from Horsham, Victoria, Australia (GPS: -36.72S, 142.18E). This clone has previously been found to lack naturally occurring facultative endosymbionts (17). We generated a *D. noxia* strain carrying *Rickettsiella* derived from *A. pisum* and another *D. noxia* strain carrying *Regiella* transinfected from *M. persicae* (see *“*Endosymbiont transinfection through microinjection”). The same clone of *D. noxia* without facultative endosymbionts (wild type) was used as a control in all experiments.

### Endosymbiont transinfection through microinjection

We introduced Rickettsiella from A. pisum and Regiella from M. persicae into apterous adult D. noxia through hemolymph microinjection (31, 79). We injected 50 D. noxia with hemolymph from A. pisum and 30 D. noxia from M. persicae, and then followed a selection step as described in Gu et al. (31) for three generations. Aphids testing positive for infection were then maintained in groups from G4 onwards. Endosymbiont status was routinely screened as well as immediately before experiments commenced through quantitative PCR (qPCR, see “Endosymbiont detection and quantification”).

### Endosymbiont detection and quantification

qPCR assays were used to confirm the presence or absence of *Rickettsiella, Regiella* and/or *Buchnera* and to measure their relative densities to a host gene following established methods (80). In the experiments assessing life history traits and population dynamics, aphids with a *Rickettsiella* and/or *Regiella* Cp value ≤ 30 and Tm values within the range of positive controls (87.5-88.3) were considered positive with high density, while aphids with a Cp value >30 were considered positive but with low density, and negative if Cp values were absent and Tm values were not within the range of positive controls. In the mesocosm experiment assessing aphid dispersal and feeding damage, aphids with a *Rickettsiella* Cp value >30 were considered *Rickettsiella* negative due to the potential for horizontally transmitted *Rickettsiella* resulting in wild type aphids testing positive with low densities (see Figure 3-figure supplement 3).

### Feeding damage to wheat and barley

*D. noxia* causes yield loss through direct feeding and leaf chlorosis by injecting toxins into the plant (81). We examined the effect of *Rickettsiella* and *Regiella* on plant damage severity to wheat and barley plants. Wheat and barley were used at Zadok’s growth stage GS14 (∼ 5 weeks old). Five 10-day-old apterous females were placed onto the second true leaf of each plant.

For wheat, individual plants were enclosed in perforated bread-stick plastic bags, secured with rubber bands and placed in randomized positions in a temperature-controlled cabinet at 25 (± 1)°C with a 16:8 h L:D photoperiod under plant growth lights. Ten replicates were established per treatment: *Rickettsiella*, *Regiella*, wild type, and no-aphid controls (40 plants in total). For barley, individual plants were placed in BugDorm cages (30 x 30 x 62 cm, mesh 160 µm aperture, Australian Entomological Supplies, Australia) and then placed in randomized positions in a temperature-controlled cabinet at 25 (± 1)°C with a 16:8h L:D photoperiod under plant growth lights. Eight replicates were established per treatment (32 plants in total). Initial aphid survival was assessed at 24 h and 48 h post-introduction in both experiments.

For each replicate plant, we recorded the number of tillers, the number of true leaves, leaf area (summed across leaves as width*length*0.75 (82)), and the percentage of leaves with chlorotic streaking. We also measured the level of aphid infestation on each plant and scored an overall plant damage score (see Supplementary File 1 for details). These assessments were undertaken either once or twice per week. All scores were assessed independently and blindly by two observers and then averaged to obtain an overall score for each plant at each time point. If the scores assigned by the two observers differed by more than one unit for a given plant, a third observer independently assessed the plant, and the average score from all three observers was used for the final score. The experiments were terminated when plants in one of the replicates had an overall plant damage score of 4 or above (Week 3.5 for wheat and Week 3 for barley). By Week 3.5, all replicate wheat plants infested with *Rickettsiella* aphids were nearly dead, with only a few aphids remaining alive. Therefore, aphid morph composition was assessed only in barley at Week 3. For this assessment, we counted the total number of nymphs, alate adults, and apterous adults on each barley plant.

### Plant defense responses

To determine the response of plants to aphid feeding with different endosymbionts, sixteen wheat plants were grown for 5 weeks as described above (see “Aphid strains and maintenance”). Four replicate plants were established for each treatment: *Rickettsiella*, *Regiella*, wild type, and no-aphid controls. For each plant, 60 aphids of mixed age (collected at random from wheat plants within eight BugDorm cages used to maintain aphids) were divided into two plastic vials (top diameter: 25 mm; height: 95 mm, DrosoKing, Australia). The top and bottom leaves of each plant were inserted into a vial and a sponge plug was placed into the vial opening to prevent aphid movement (see Figure 2A). Control plants were treated in same manner, except no aphids were added to the vials. All plants were then placed in randomized positions in a temperature-controlled cabinet at 25 (± 1)°C with a 16:8 h L:D photoperiod under plant growth lights. After a 7-day infestation period, all aphids were removed and the plants maintained for an additional 7 days. This time period was chosen based on a pilot experiment indicating that endogenous JA and SA typically display distinct changes during this time.

For the sample collection, we combined the upper and lower leaves together to provide a more representative measure of the plant-level response, rather than focusing on responses at a particular leaf position. Plant material (∼200 mg) was excised from the top and bottom leaves (aphid feeding areas) and the middle leaf (non-feeding area) of each wheat plant was placed in 60 mL centrifuge tubes and immediately frozen in liquid nitrogen. Fifty mg of this material was then cryomilled (3 x 30 sec, 3620 *g*) in 600 µL of 3:1 MeOH/water containing 0.28 nmol of ^13^C_5_ ^15^N valine, ^13^C_5_ sorbitol, dihydrojasmonic acid and salicylic acid-d6, using a

Bertin Technologies Precellys bead mill coupled to a Cryolys cooling unit. 450 µL of each homogenate was transferred into fresh Eppendorf tubes and centrifuged at 4°C for 10 min at 16,000 *g* using an Eppendorf centrifuge 5430 R. The resulting supernatants were then transferred into fresh 1.5 mL Eppendorf tubes. An 80 µL aliquot of each sample was pooled to create the pooled biological quality control (PBQC). Two hundred µL of each study sample and the PBQC were transferred into HPLC inserts and evaporated at 30°C to complete dryness, using a CHRIST RVC 2-33 CD plus speed vacuum. To limit the amount of moisture present in the insert, 50 µL 100% methanol (LCMS grade) was added to each insert and evaporated using a speed vacuum. Dried samples for GCMS analysis were derivatised online using the Shimadzu AOC6000 autosampler robot. Derivatisation was achieved by the addition of 25 µL methoxyamine hydrochloride (30 mg/mL in pyridine, Merck) followed by shaking at 37°C for 2h. Samples were then derivatised with 25 µL of N,O-bis (trimethylsilyl)trifluoroacetamide with trimethylchlorosilane (BSTFA with 1% TMCS, Thermo Scientific) for 1h at 37°C. The sample was allowed to equilibrate at room temperature for 1 h before 1 µL was injected onto the GC column using a hot needle technique. A splitless injection was performed for each sample.

The GC-MS system used comprised of an AOC6000 autosampler, a 2030 Shimadzu gas chromatograph and a TQ8050NX triple quadrupole mass spectrometer (Shimadzu, Japan) with an electron ionisation source (-70eV). The mass spectrometer was tuned according to the manufacturer’s recommendations using tris-(perfluorobutyl)-amine (CF43). GC-MS was performed on a 30m Agilent DB-5 column with 0.25mm internal diameter column and 1µm film thickness. The injection temperature (inlet) was set at 280°C, the MS transfer line at 280°C and the ion source adjusted to 200°C. Helium was used as the carrier gas at a flow rate of 1 mL/min and argon gas was used in the collision cell to generate the MRM product ion. The analysis of the derivatised samples was performed under the following oven temperature program; 100°C start temperature, hold for 4 minutes, followed by a 10°C/min oven temperature ramp to 320°C with a following final hold for 11 minutes. Samples were analysed in a randomised order, with a pooled biological quality control sample added after every 5 samples. The pooled biological quality control was used to monitor downstream sample stability and analytical reproducibility. Approximately 625 targets were collected using the Shimadzu Smart Metabolite Database, where each target comprised a quantifier MRM along with a qualifier MRM, which covers approximately 420 unique endogenous metabolites and multiple stable isotopically labelled internal standards. Resultant data was processed using Shimadzu LabSolutions Insight software, where peak integrations were visually validated and manually corrected where required. JA and JA-Ile, were quantified by comparing the peak area to the spike in dihydrojasmonic acid (not endogenously found) and SA was quantified by comparing the peak area to the spike in salicylic acid-d6 standard. All the other components were normalized by the peak area to the spike in the ^13^C_5_ ^15^N valine.

### Aphid life history traits at two temperatures

We measured the effects of *Rickettsiella* and *Regiella* on development time, lifetime fecundity and longevity following the methods described by Gu et al. (31). Two nymphs (< 24 h old) were transferred to a plastic cup (150 ml: 8 x 6 x 5 cm, Daily Best, Auburn, Australia) containing and a single wheat plant (GS14, with the root removed) inserted into 1% agar. A second cup was placed on top and sealed with Parafilm. Sixty replicates were established per treatment: *Rickettsiella*, *Regiella* and wild type, with thirty cups randomly positioned in a temperature-controlled cabinet at 19 (± 1)°C and the remaining thirty randomly positioned in a temperature-controlled cabinet at 25 (± 1)°C. Both cabinets were programmed with a 16:8h L:D photoperiod under plant growth lights. After 7 d, a single aphid was randomly removed and stored in 100% ethanol and frozen for later endosymbiont density measurements. The remaining aphid from each cup was transferred to new cups weekly to assess body color and body shape, fecundity and longevity. The *Rickettsiella* and *Regiella* strains were compared to the wild type in separate experiments, allowing for more replicates to be included.

### Body color and body size

To measure the effects of *Regiella* on body color and body size, we removed apterous adults (∼ 14 d old) from individual cups containing wheat plants. The aphids that were maintained at both 19 (± 1)°C and 25 (± 1)°C in temperature-controlled cabinets with a 16:8h L:D photoperiod under plant growth lights. We then measured body length and body color by taking photos and analyzing these with ImageJ, as described previously (31). We assessed 25 individuals per strain: *Regiella* and wild type. Different to effects seen in other aphid species (e.g. 15, 31, 41), there was no obvious effect of *Rickettsiella* on body color after transinfection to *D. noxia*. Nevertheless, to test for subtle differences, we repeated the body color methods above for *Rickettsiella* and wild type aphids, except that we only examined aphids at 19 (± 1)°C.

### Population dynamics in mixed cages

We assessed the population dynamics of *Rickettsiella* and *Regiella* over multiple generations at two different temperatures. We mixed 30 wild type apterous adults (∼ 14 d old) with 30 *Rickettsiella* or 30 *Regiella* apterous adults (∼ 14 d old) and transferred these to six wheat plants (∼ 6 weeks old), which were placed inside a single BugDorm cage (32.5 x 32.5 x 77 cm, mesh 160 µm aperture, Australian Entomological Supplies, Australia). Each cage was replicated five times and these were placed in randomized positions in a temperature-controlled cabinet at 19 (± 1)°C with a 16:8h L:D photoperiod under plant growth lights. These experiments were repeated under the same conditions, except the cages were placed at 25 (± 1)°C. Aphids were transferred to new wheat plants every 3 weeks at 19°C and every 2 weeks at 25°C. During each transfer, 60 randomly selected aphids were placed onto the new plants, with the remaining aphids stored in 100% ethanol and frozen for later endosymbiont density measurements. For *Regiella,* the 19°C experiment ran for 24 weeks and the 25°C experiment ran for 30 weeks. The *Rickettsiella* experiments ran for 12 weeks for both temperatures. To measure *Rickettsiella* and *Regiella* frequency over time, we screened 15-20 aphids per replicate cage at multiple time points using qPCR (see “Endosymbiont detection and quantification*”).*

### Population growth and alate production on whole plants

Differences in aphid morphs were observed on barley in the feeding damage experiment, and here we measured aphid population growth and alate production on whole wheat plants. The measurements for *Rickettsiella* and *Regiella* aphids were conducted separately. Different methods were used for the two endosymbionts to reflect specific research aims based on earlier data on aphid morphs and the severity of feeding damage to plants. For *Rickettsiella*, the primary focus was on alate frequency, which was observed to increase at later stages when plants were under stress in the “Feeding damage to barley” experiment. In contrast, for *Regiella*, the emphasis was on early population growth and aphid distribution across the plant due to the potential impact of *Regiella* on reducing feeding damage to plants.

For the *Rickettsiella* experiment, five 14-day-old apterous females were placed onto the second true leaf of a single wheat plant (GS14) (see Figure 3A). Each plant was placed into a BugDorm cage (32.5 x 32.5 x 77 cm, mesh 160 µm aperture) and placed in a randomized position in a temperature-controlled cabinet at 25 (± 1)°C with a 16:8h L:D photoperiod under plant growth lights. Nine replicates were established per treatment: *Rickettsiella* and wild type. Initial survival of aphids was assessed after 24 h and 48 h. At day 14, three replicate plants were randomly selected from the available plants for both treatments and we counted the total number of nymphs, alate adults, and apterous adults. This was repeated again at Days 21 and 25.

For the *Regiella* experiment, five 14-day-old apterous females were placed onto each of the top, middle and bottom leaves of a single wheat plant (GS14). A plastic vial was placed over each leaf and cotton wool inserted into the vial opening to prevent aphid movement (see Figure 4A). Each plant was placed into a BugDorm cage (32.5 x 32.5 x 77 cm, mesh 160 µm aperture) and placed in a randomized position in a temperature-controlled cabinet at 25 (± 1)°C with a 16:8h L:D photoperiod under plant growth lights. Sixteen replicates for wild type and seventeen replicates for *Regiella* were established. Initial survival of aphids was again assessed after 24 h and 48 h. Four plants per treatment were originally planned for sampling at each time point (Days 4, 8, 12, and 16). However, because clear differences in plant damage were observed between treatments by Day 12, one additional plant per treatment was measured at Day 12 (*n* = 5). The remaining plants were measured at Day 16.

### Aphid dispersal ability and plant feeding damage in mesocosms

Given the number of alate and apterous adults in our feeding damage experiment on individual plants varied among aphid strains (Figure 1-figure supplement 1I & Figure 3C), we undertook a larger-scale experiment to test whether the reduced alate production observed with *Rickettsiella* also translated into reduced dispersal at the population level. We did not conduct the same experiment with *Regiella* because we did not detect a difference in alate frequency in our earlier experiments (Figure 1-figure supplement 1I & Figure 4C). To do this, three plastic trays (39.5 x 28 x 11cm, Bargain Basement, Australia), each containing eight individual wheat plants (∼5-week-old, GS14), were established. These trays were placed inside a large BugDorm cage (93 x 47.5 x 47.5cm, mesh 160 µm aperture), creating a mesocosm environment. Five replicate mesocosms were established and randomly positioned in a temperature-controlled room at 25 (± 1)°C with a 16:8 h L:D photoperiod under plant growth lights. In each mesocosm, *Rickettsiella* and wild type aphids were released at either end of the BugDorm cages, with five 14-day-old apterous females placed on the second true leaf of each wheat plant (*n* = 4 wheat plants each side) (see Figure 5A). The four plants with aphids represented the ‘aphid-release’ wheat and the four adjacent plants represented the ‘aphid-spread’ wheat. The middle tray with wheat plants was used to monitor aphid dispersal.

Initial aphid survival was assessed after 24 h and 48 h, and the experiment ran for 5 weeks. Once or twice per week, we recorded the total number of leaves, leaf area, the percentage of leaves with chlorotic streaking, aphid colonization rate, the level of aphid infestation and scored an overall plant damage score (see Supplementary File 1) for all ‘aphid-release’ and ‘aphid-spread’ plants (total 16 plants per replicate). All scores were assessed independently and blindly by two observers and then averaged to obtain an overall score for each plant at each time point. If the scores assigned by the two observers differed by more than one unit for a given plant, a third observer independently assessed the plant, and the average score from all three observers was used for the final score. To assess aphid dispersal, all eight wheat plants in the middle ‘dispersal’ tray were removed weekly and replaced with fresh plants at the same plant growth stage. All aphids on these eight plants were subsequently removed, and the number of alate and apterous individuals counted. In addition, 20 alates and 20 apterous aphids at Weeks 2 and 3 within each mesocosm were randomly removed, placed into 100% ethanol and later screened for *Ricketsiella* infection.

### Statistical analysis

Life history traits on small plants, aphid population data from large cages, and aphid feeding damage data were analysed using IBM SPSS Statistics 29.0 for Mac following tests for normality. For life history traits on small plants, development time measures were analysed with Kruskal-Wallis tests. Fecundity was analysed with general linear models (GLMs), with developmental temperature, endosymbiont status (*Rickettsiella, Regiella* and wild type) and their interaction included as factors. Endosymbiont density values were compared by independent sample t-tests. We used Cox regression to assess the impact of *Rickettsiella* and *Regiella* on aphid longevity. For body color, body shape, and body size, we analysed each component (hue, saturation, lightness) separately using independent sample t-tests as well as all the components together as a multivariate ANOVA (MANOVA). For the population growth experiment in large cages, GLMs were used to test overall patterns in aphid numbers across relevant time periods, including *Rickettsiella* (Days 21 and 25) and *Regiella* (Days 8–16) aphid strains. Sampling date was log-transformed and normalised prior to analysis. The independent sample t-test was used to compare differences between endosymbiont and wild type treatments at each time point. The earliest time points in the experiments for *Rickettsiella* (Day 14) and *Regiella* (Day 4) were excluded because aphid numbers were similar across all developmental stages at the start of the experiment. For the feeding damage experiments, we used repeated measures in GLMs for the wheat and barley feeding damage experiment and the mesocosm wheat experiment. Aphid dispersal data from the mesocosm experiment was analysed by paired t-tests. Where more than two treatments were compared, we used probabilities from two treatment comparisons (after Bonferroni correction) to determine significance.

All statistical analyses and visualisations for plant responses were conducted in R studio (version 4.5.2) for Mac. Metabolomic data from wheat plant tissues were normalised to the internal standard valine prior to analysis to correct for technical variation. The dataset was normalised by median and log transformed. Principal component analysis (PCA) was performed using all detected components to visualise overall metabolic variation among treatments. Heatmaps were generated using the top 50 components with the highest variance across samples. Differences in all the component abundance between treatments were assessed using linear or generalized linear models, and components with *p* < 0.05 were considered significantly different.

All figures were generated using Prism 10 for Mac and R studio with the ggplot2 and pheatmap packages.

## Acknowledgements

We thank David de Souza, Jordi Hondrogiannis, Courtney Brown, Ella Yeatman and Kelly Richardson for technical assistance, the Grains Innovative Park for supplying the *D. noxia* colony. This research used NCRIS-enabled Metabolomics Australia infrastructure at the University of Melbourne and funded through BioPlatforms Australia. This work was undertaken as part of the Australian Grains and Horticulture Pest Innovation Program (AGHPIP), supported through funding provided by the Grains Research and Development Corporation (UOM1906-002RTX; UOM2404-006RT), and by Hort Innovation Australia (ST23002) with additional support from the University of Melbourne and Cesar Australia.

Materials and data availability: The aphid lines generated in this study are available from the corresponding author upon reasonable request. All data underlying the results presented in this study are provided in the source raw data files associated with the manuscript.

