## Supplementary figures and images for "Novel but stable endosymbionts have contrasting effects on aphid dispersal and plant feeding damage in the cereal pest *Diuraphis noxia*"

### Fig 1-S1

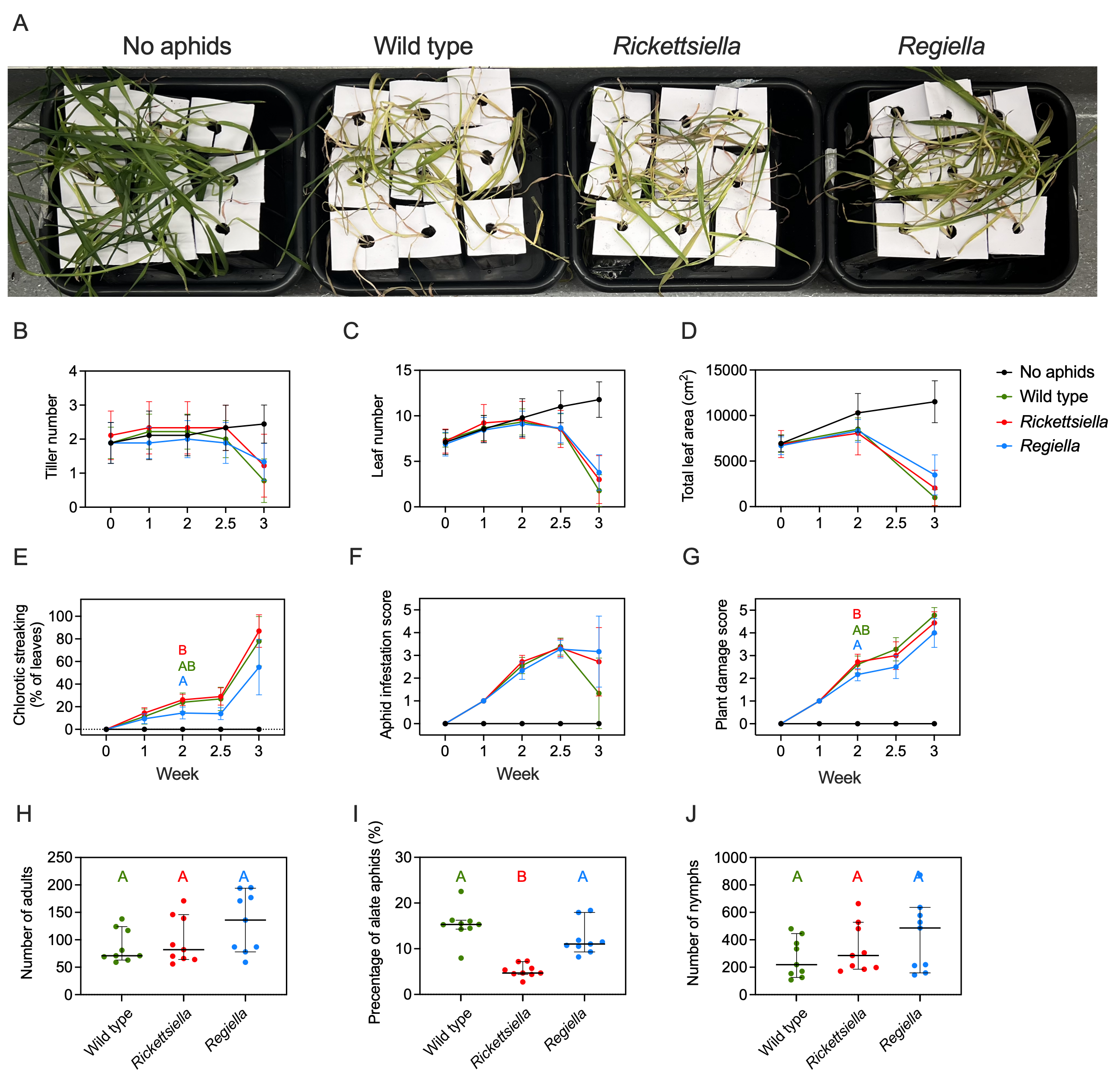

### Fig 2-S1

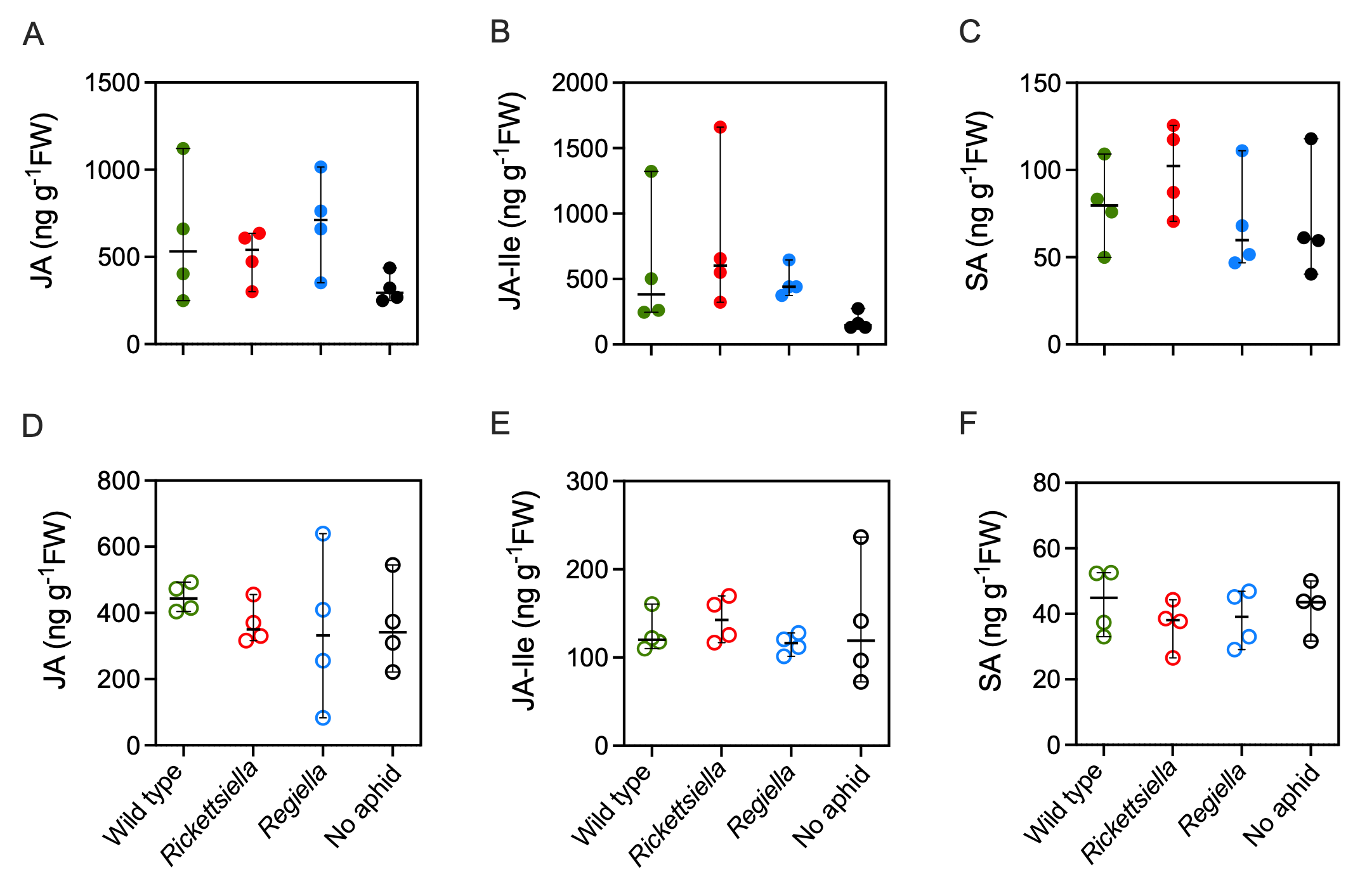

### Fig 2-S2

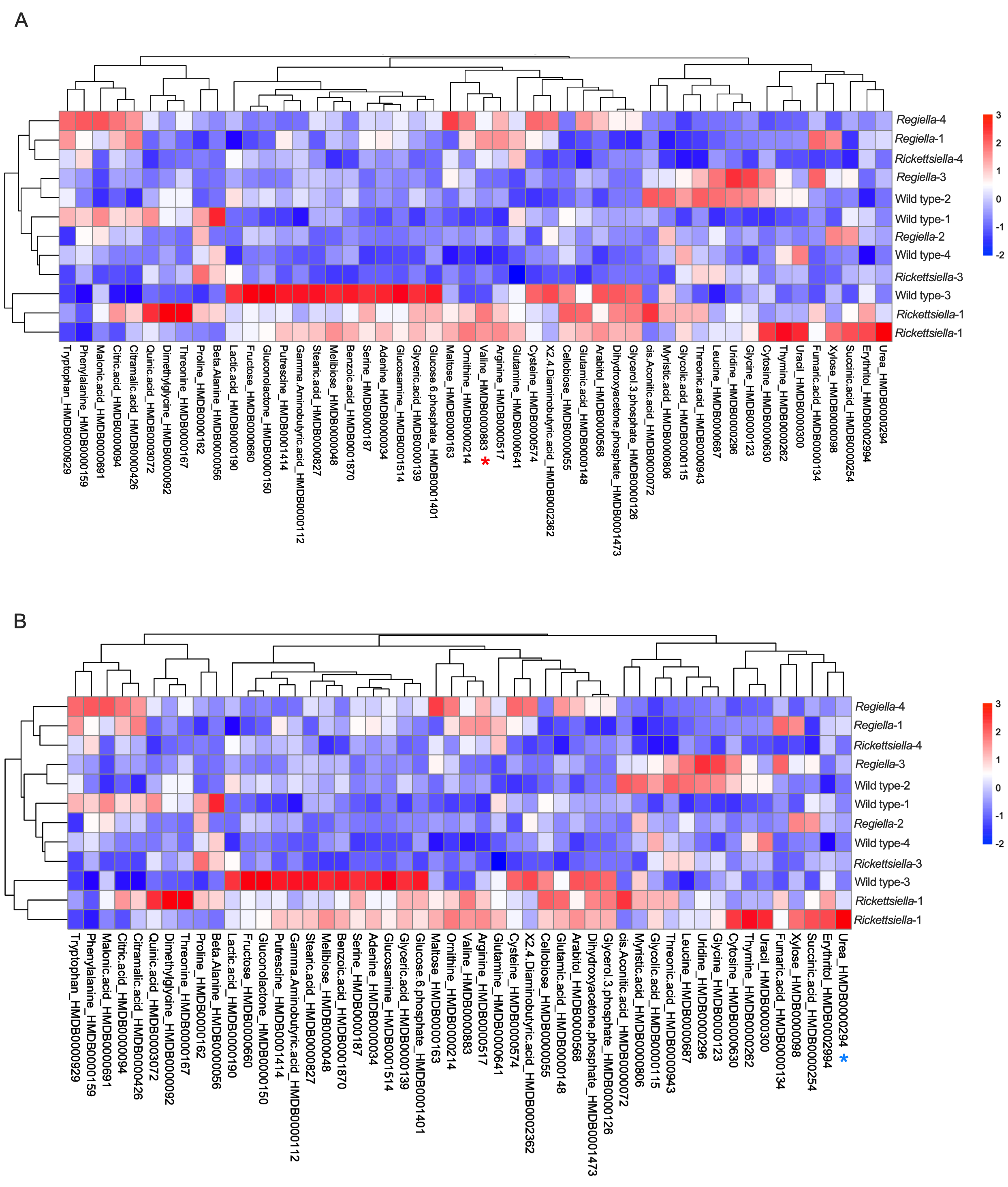

### Fig 2-S3

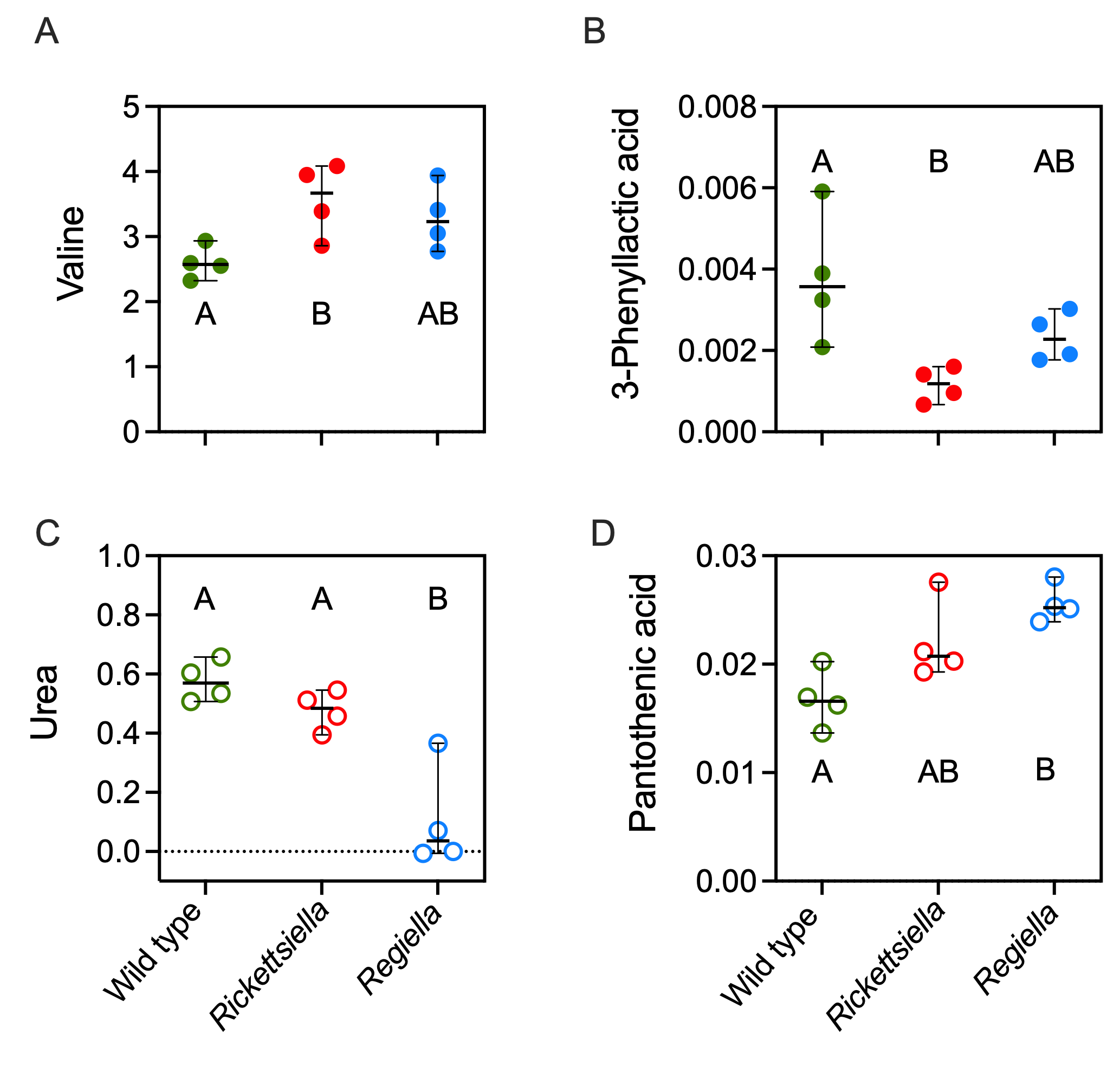

### Fig 3-S1

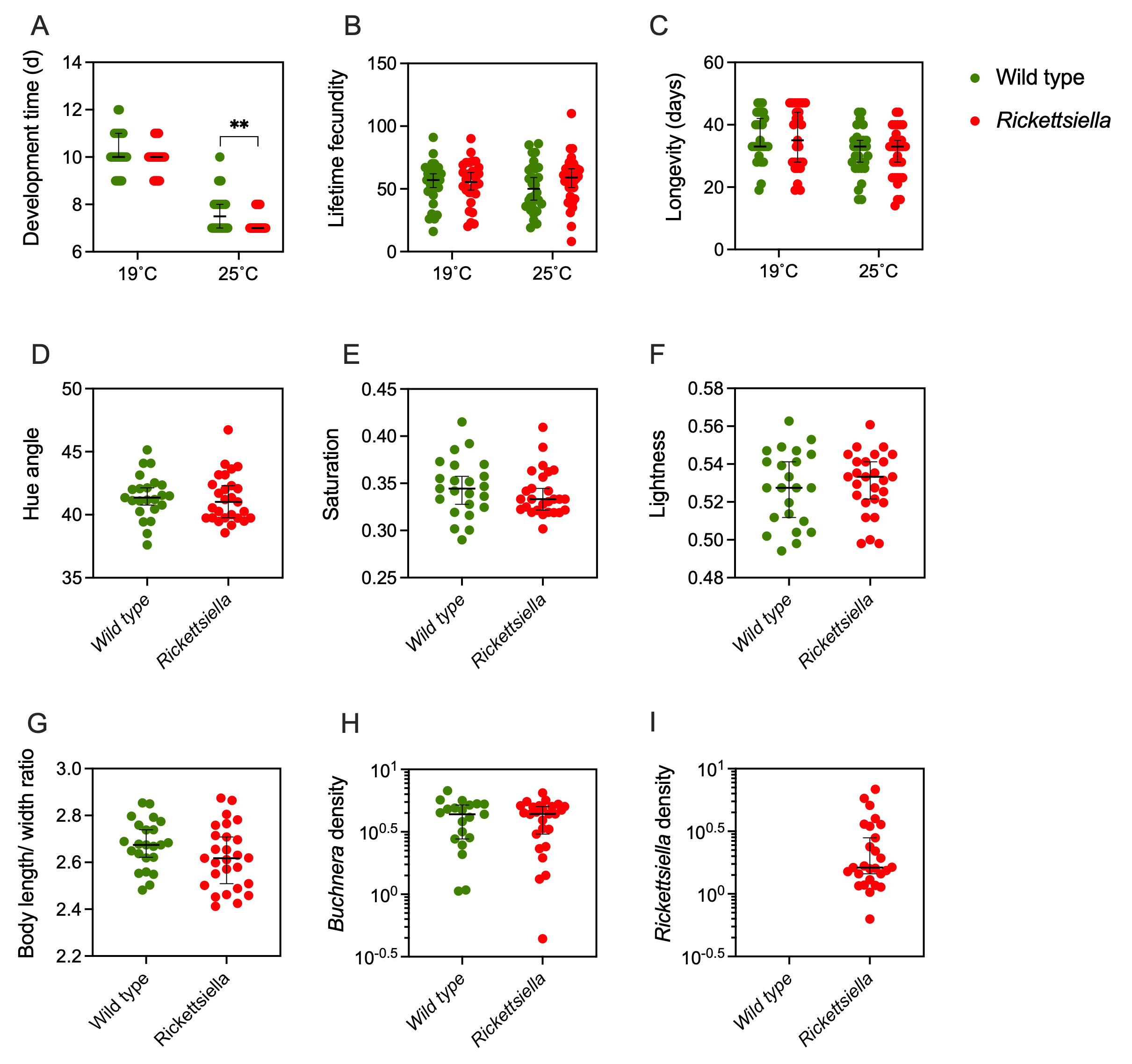

### Fig 3-S2

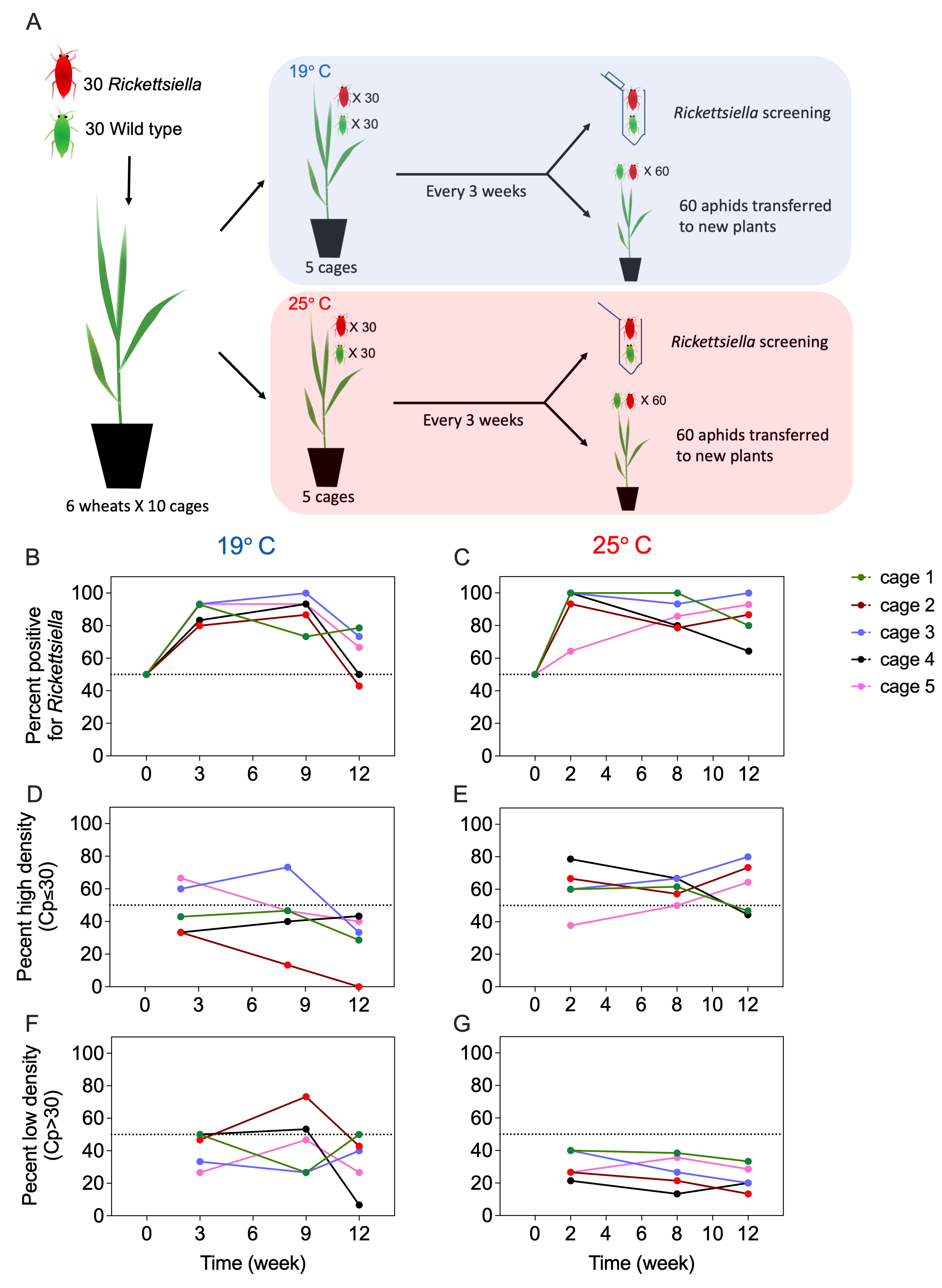

### Fig 3-S3

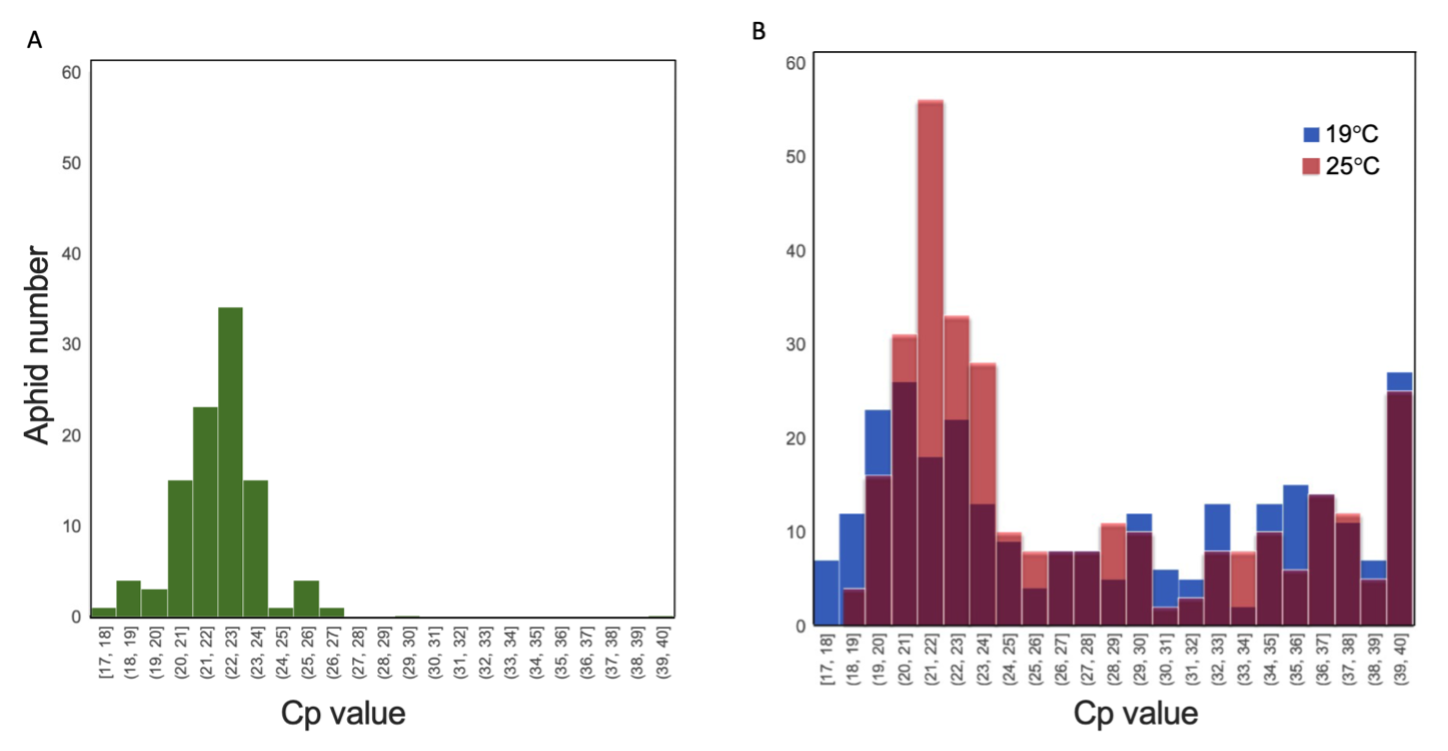

### Fig 4-S1

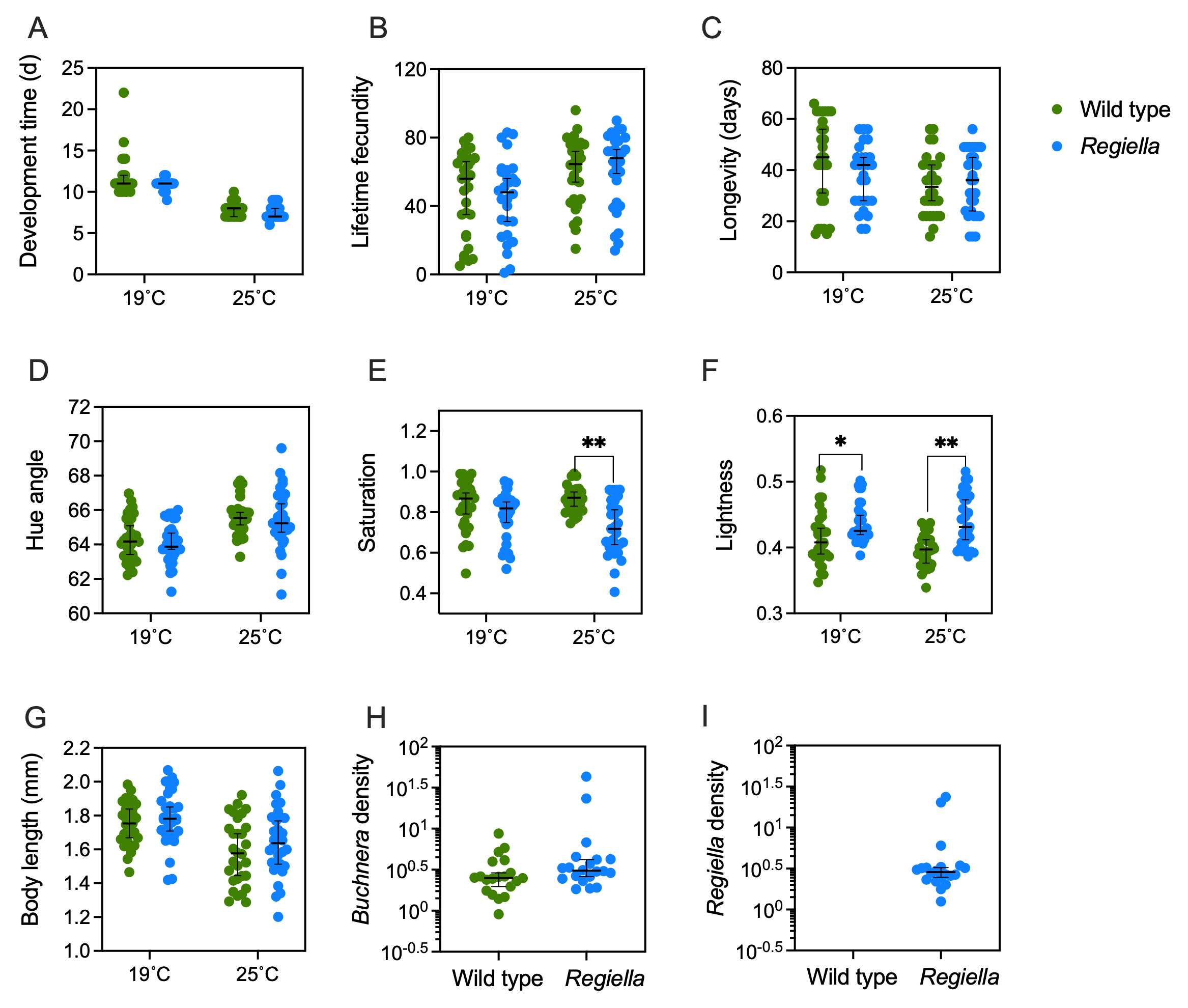

### Fig 4-S2

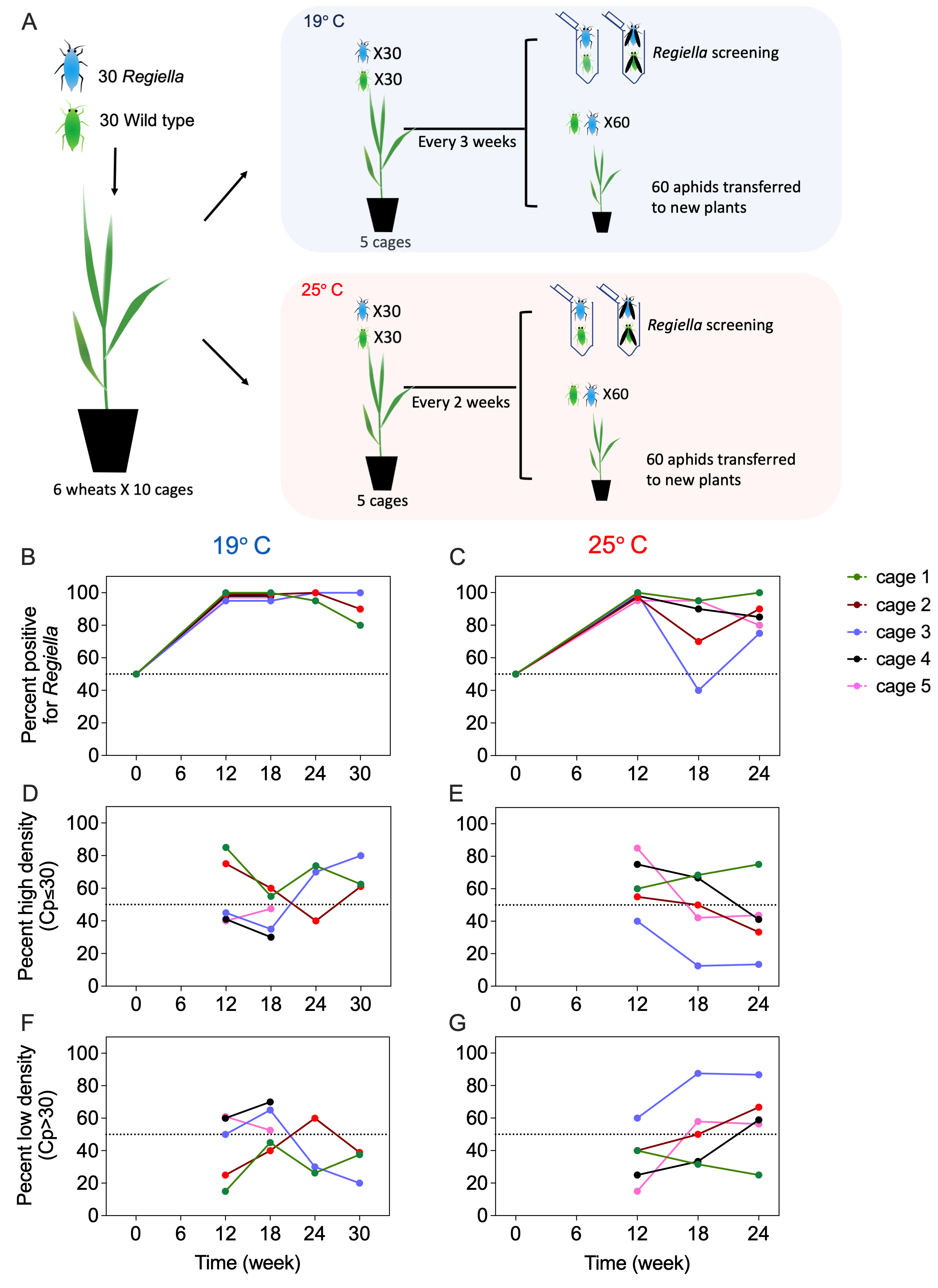

### Fig 4-S3

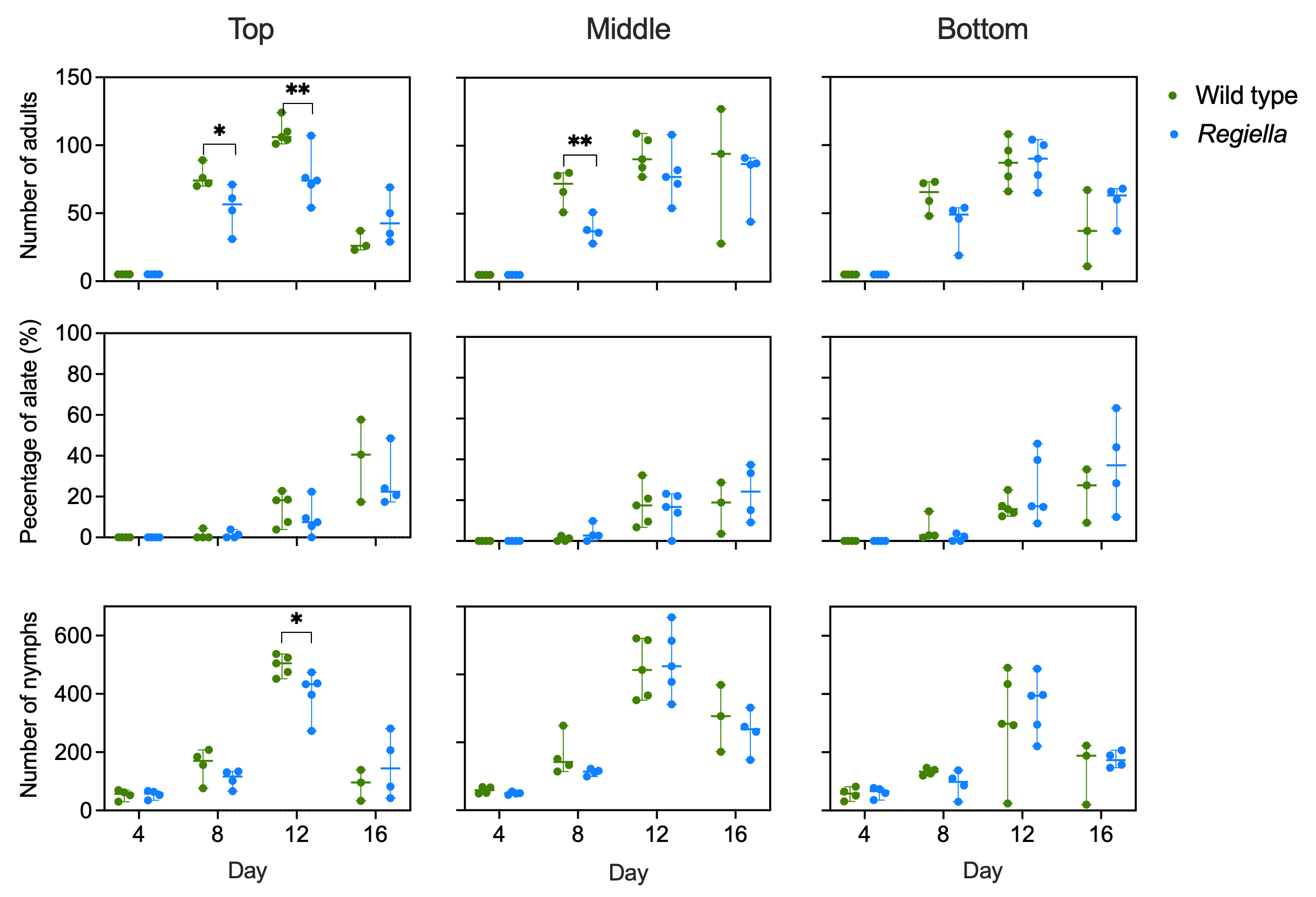
