## Supplemental document for "Novel but stable endosymbionts have contrasting effects on aphid dispersal and plant feeding damage in the cereal pest *Diuraphis noxia*"

Short title: Endosymbionts impact aphid dispersal and plant feeding damage

Xinyue Gu^1^, Alex Gill^1^, Qiong Yang^1^, Perran A. Ross^1,2^, Lucy Hayward^1^, Monica Stelmach^1^, Paul A. Umina^1,3^, Sheik Nadeem Elahee Doomun^4^, Mel Berran^1^, Lauren Coakley^1^, Sonia Sharma^1^, Ary A. Hoffmann^1^

^1^Pest and Environmental Adaptation Research Group, School of BioSciences, Bio21 Molecular Science and Biotechnology Institute, The University of Melbourne, Parkville, Australia

^2^Section for Bioscience and Engineering, Department of Chemistry and Bioscience, Aalborg University, Aalborg, Denmark

^3^Cesar Australia, Brunswick, Australia

^4^Metabolomics Australia, Bio21 Molecular Science and Biotechnology Institute, The University of Melbourne, Parkville, Australia

**Classification:** Applied Biological Sciences, microbiology

### ****References****

1. Kikuchi Y. 2009. Endosymbiotic bacteria in insects: their diversity and culturability. Microbes Environ 24:195–204.
2. Moran NA, McCutcheon JP, Nakabachi A. 2008. Genomics and evolution of heritable bacterial symbionts. Annu Rev Genet 42:165–190.
3. Duron O et al. 2008. The diversity of reproductive parasites among arthropods: *Wolbachia* do not walk alone. BMC Biol 6:27.
4. Michalkova V, Benoit JB, Weiss BL, Attardo GM, Aksoy S. 2014. Vitamin B6 generated by obligate symbionts is critical for maintaining proline homeostasis and fecundity in tsetse flies. Appl Environ Microbiol 80:5844–5853.
5. Douglas AE. 1998. Nutritional interactions in insect-microbial symbioses: aphids and their symbiotic bacteria Buchnera. Annu Rev Entomol 43:17–37.
6. Stillson PT, Martinez K, Adamson J, Tehrani A, Ravenscraft A. 2025. Temperature influences outcomes of an environmentally acquired symbiosis. ISME J 19:wraf056.
7. Gu XY et al. 2022. A wMel Wolbachia variant in Aedes aegypti from field-collected Drosophila melanogaster with increased phenotypic stability under heat stress. Environ Microbiol 24:2119–2135.
8. Zhang B, Leonard SP, Li YY, Moran NA. 2019. Obligate bacterial endosymbionts limit thermal tolerance of insect host species. Proc Natl Acad Sci USA 116:24712–24718.
9. Stillson PT, Sim SB, Corpuz RL, Ravenscraft A. 2025. Symbiont gene expression predicts insect host's response to high temperatures. Mol Ecol 34:e70154.
10. Pais R, Lohs C, Wu YN, Wang JW, Aksoy S. 2008. The obligate mutualist Wigglesworthia glossinidia influences reproduction, digestion, and immunity processes of its host, the tsetse fly. Appl Environ Microbiol 74:5965–5974.
11. Rosenwald LC, Sitvarin MI, White JA. 2020. Endosymbiotic Rickettsiella causes cytoplasmic incompatibility in a spider host. Proc R Soc B 287:20201107.
12. Zabalou S et al. 2004. Wolbachia-induced cytoplasmic incompatibility as a means for insect pest population control. Proc Natl Acad Sci USA 101:15042–15045.
13. Hrdina A, Canales MS, Arias-Rojas A, Frahm D, Iatsenko I. 2024. The endosymbiont Spiroplasma poulsonii in Drosophila melanogaster increases resistance to pathogens by enhancing iron sequestration and melanization. mBio 15:e00936-24.
14. Massey JH, Newton ILG. 2022. Diversity and function of arthropod endosymbiont toxins. Trends Microbiol30:185–198.
15. Yu WJ et al. 2025. A persistent bacterial Regiella transinfection in the bird cherry-oat aphid Rhopalosiphum padi increasing host fitness and decreasing plant virus transmission. Pest Manag Sci 81:2791–2799.
16. Shi PQ et al. 2024. Rickettsia transmission from whitefly to plants benefits herbivore insects but is detrimental to fungal and viral pathogens. mBio 15:e02448-23.
17. Yang Q et al. 2023. A diversity of endosymbionts across Australian aphids and their persistence in aphid cultures. Environ Microbiol 25:1988–2001.
18. Guyomar C et al. 2018. Multi-scale characterization of symbiont diversity in the pea aphid complex through metagenomic approaches. Microbiome 6:181.
19. Fukatsu T. 1994. Endosymbiosis of aphids with microorganisms: a model case of dynamic endosymbiotic evolution. Plant Species Biol 9:145–154.
20. Vorburger C, Gehrer L, Rodriguez P. 2010. A strain of the bacterial symbiont Regiella insecticola protects aphids against parasitoids. Biol Lett 6:109–111.
21. Tsuchida T, Koga R, Matsumoto S, Fukatsu T. 2011. Interspecific symbiont transfection confers a novel ecological trait to the recipient insect. Biol Lett 7:245–248.
22. Kang ZW et al. 2022. Facultative endosymbiont Serratia symbiotica inhibits the apterization of pea aphid to enhance its spread. Microbiol Spectr 10:e04066-22.
23. Hansen AK, Moran NA. 2014. The impact of microbial symbionts on host plant utilization by herbivorous insects. Mol Ecol 23:1473–1496.
24. Dedryver CA, Le Ralec A, Fabre F. 2010. The conflicting relationships between aphids and men: a review of aphid damage and control strategies. C R Biol 333:539–553.
25. Valenzuela I, Hoffmann AA. 2015. Effects of aphid feeding and associated virus injury on grain crops in Australia. Austral Entomol 54:292–305.
26. Tatchell GM. 1989. An estimate of the potential economic losses to some crops due to aphids in Britain. Crop Prot 8:25–29.
27. Edwards OR, Franzmann B, Thackray D, Micic S. 2008. Insecticide resistance and implications for future aphid management in Australian grains and pastures: a review. Aust J Exp Agric 48:1523–1530.
28. Reyes ML et al. 2019. The influence of symbiotic bacteria on reproductive strategies and wing polyphenism in pea aphids responding to stress. J Anim Ecol 88:601–611.
29. Sanches P, De Moraes CM, Mescher MC. 2023. Endosymbionts modulate virus effects on aphid-plant interactions. ISME J 17:2441–2451.
30. Gu XY et al. 2024. Influence of genetic and environmental factors on the success of endosymbiont transfers in pest aphids. Environ Microbiol 26:e16704.
31. Gu XY et al. 2023. A rapidly spreading deleterious aphid endosymbiont that uses horizontal as well as vertical transmission. Proc Natl Acad Sci USA 120:e2217278120.
32. Gu XY et al. 2025. Transinfections of the endosymbiont Rickettsiella viridis in different Myzus persicae (Hemiptera: Aphididae) clones show consistent deleterious effects and stable transmission. J Econ Entomol118:1544–1552.
33. Ross PA et al. 2024. Deleterious effects of the endosymbiont Rickettsiella viridis in Myzus persicae are environmentally dependent. J Pest Sci 98:375–388.
34. Luna E et al. 2018. Bacteria associated with Russian wheat aphid (Diuraphis noxia) enhance aphid virulence to wheat. Phytobiomes J 2:151–164.
35. Summers CG, Newton AS, Kirk M, Temple SR. 1990. Transmission of beet yellows and beet mosaic viruses by noncolonizing aphid vectors. J Econ Entomol 83:2448–2451.
36. Halbert SE, Mink GI, Silbernagel MJ, Mowry TM. 1994. Transmission of bean common mosaic virus by cereal aphids (Homoptera: Aphididae). Plant Dis 78:983–985.
37. Ennahli S et al. 2009. Comparison of categories of resistance in wheat and barley genotypes against biotype 2 of the Russian wheat aphid, Diuraphis noxia (Kurdjumov). Arthropod Plant Interact 3:45–53.
38. Stoetzel MB. 1987. Information on and identification of Diuraphis noxia (Homoptera: Aphididae) and other aphid species colonizing leaves of wheat and barley in the United States. J Econ Entomol 80:696–704.
39. Weiland AA et al. 2008. Biotypic diversity in Colorado Russian wheat aphid (Hemiptera: Aphididae) populations. J Econ Entomol 101:569–574.
40. Puterka GJ et al. 2015. Change in biotypic diversity of Russian wheat aphid (Hemiptera: Aphididae) populations in the United States. J Econ Entomol 108:798–804.
41. Tsuchida T et al. 2010. Symbiotic bacterium modifies aphid body color. Science 330:1102–1104.
42. Botha AM, Li YC, Lapitan NLV. 2005. Cereal host interactions with Russian wheat aphid: a review. J Plant Interact 1:211–222.
43. Vierstra RD. 1996. Proteolysis in plants: mechanisms and functions. Plant Mol Biol 32:275–302.
44. Malatrasi M et al. 2006. A branched-chain amino acid aminotransferase gene isolated from Hordeum vulgare is differentially regulated by drought stress. Theor Appl Genet 113:965–976.
45. Aidoo MK et al. 2017. A combination of stomata deregulation and a distinctive modulation of amino acid metabolism are associated with enhanced tolerance of wheat varieties to transient drought. Metabolomics 13:138.
46. Mu WM, Yu SH, Zhu LJ, Zhang T, Jiang B. 2012. Recent research on 3-phenyllactic acid, a broad-spectrum antimicrobial compound. Appl Microbiol Biotechnol 95:1155–1163.
47. Witte CP. 2011. Urea metabolism in plants. Plant Sci 180:431–438.
48. Coxon KM et al. 2005. Pantothenate biosynthesis in higher plants. Biochem Soc Trans 33:743–746.
49. Golan K et al. 2017. Accumulation of amino acids and phenolic compounds in biochemical plant responses to feeding of two different herbivorous arthropod pests. Arthropod Plant Interact 11:675–682.
50. Divekar PA et al. 2022. Plant secondary metabolites as defense tools against herbivores for sustainable crop protection. Int J Mol Sci 23:2690.
51. Li Q, Fan J, Sun JX, Wang MQ, Chen JL. 2018. Plant-mediated horizontal transmission of Hamiltonella defensa in the wheat aphid Sitobion miscanthi. J Agric Food Chem 66:13367–13377.
52. Pons I, Renoz F, Noël C, Hance T. 2019. Circulation of the cultivable symbiont Serratia symbiotica in aphids is mediated by plants. Front Microbiol 10:764.
53. Pons I, Renoz F, Noël C, Hance T. 2019. New insights into the nature of symbiotic associations in aphids: infection process, biological effects, and transmission mode of cultivable Serratia symbiotica bacteria. Appl Environ Microbiol 85:e02445-02418.
54. Perreau J et al. 2021. Vertical transmission at the pathogen-symbiont interface: Serratia symbiotica and aphids. mBio 12:e00359-21.
55. Liu XD, Lei HX, Chen FF. 2019. Infection pattern and negative effects of a facultative endosymbiont on its insect host are environment-dependent. Sci Rep 9:4013.
56. Zepeda-Paulo F, Villegas C, Lavandero B. 2017. Host genotype-endosymbiont associations and their relationship with aphid parasitism at the field level. Ecol Entomol 42:86–95.
57. Bennett AE, Millar NS, Gedrovics E, Karley AJ. 2016. Plant and insect microbial symbionts alter the outcome of plant-herbivore-parasitoid interactions: implications for invaded, agricultural and natural systems. J Ecol 104:1734–1744.
58. Lukasik P, Guo H, Van Asch M, Ferrari J, Godfray HCJ. 2013. Protection against a fungal pathogen conferred by the aphid facultative endosymbionts Rickettsia and Spiroplasma is expressed in multiple host genotypes and species and is not influenced by co-infection with another symbiont. J Evol Biol 26:2654–2661.
59. Simon JC, Peccoud J. 2018. Rapid evolution of aphid pests in agricultural environments. Curr Opin Insect Sci 26:17–24.
60. Zhang FM et al. 2015. Bacterial symbionts, Buchnera, and starvation on wing dimorphism in English grain aphid, Sitobion avenae (F.) (Homoptera: Aphididae). Front Physiol 6:155.
61. Yang Q et al. 2026. A Rickettsiella transinfection in Rhopalosiphum padi reduces fitness and alate production but not plant virus transmission. Pest Manag Sci 82:3894–3906.
62. Müller CB, Williams IS, Hardie J. 2001. The role of nutrition, crowding and interspecific interactions in the development of winged aphids. Ecol Entomol 26:330–340.
63. Leybourne DJ, Valentine TA, Bos JIB, Karley AJ. 2020. A fitness cost resulting from Hamiltonella defensa infection is associated with altered probing and feeding behaviour in Rhopalosiphum padi. J Exp Biol 223:jeb207936.
64. Li XA et al. 2023. Sitobion miscanthi L type symbiont enhances the fitness and feeding behavior of the host grain aphid. Pest Manag Sci 79:1362–1371.
65. Loudit SMB, Bauwens J, Francis F. 2018. Cowpea aphid-plant interactions: endosymbionts and related salivary protein patterns. Entomol Exp Appl 166:460–473.
66. De Jager LE. 2014. Characterization of the mitochondrial genomes of Diuraphis noxia biotypes. MSc thesis, Stellenbosch University.
67. Clavé C et al. 2022. Physiological costs of facultative endosymbionts in aphids assessed from energy metabolism. Funct Ecol 36:2580–2592.
68. Chen YZ et al. 2020. An aphid RNA transcript migrates systemically within plants and is a virulence factor. Proc Natl Acad Sci USA 117:12763–12771.
69. du Preez PH et al. 2020. DNA methylation and demethylation are regulated by functional DNA methyltransferases and DnTET enzymes in Diuraphis noxia. Front Genet 11:452.
70. Puterka GJ et al. 2014. Characterization of eight Russian wheat aphid (Hemiptera: Aphididae) biotypes using two-category resistant-susceptible plant responses. J Econ Entomol 107:1274–1283.
71. Bru D, Martin-Laurent F, Philippot L. 2008. Quantification of the detrimental effect of a single primer-template mismatch by real-time PCR using the 16S rRNA gene as an example. Appl Environ Microbiol 74:1660–1663.
72. Simonet P et al. 2016. Direct flow cytometry measurements reveal a fine-tuning of symbiotic cell dynamics according to the host developmental needs in aphid symbiosis. Sci Rep 6:19967.
73. James EB, Pan X, Schwartz O, Wilson ACC. 2022. SymbiQuant: a machine learning object detection tool for polyploid independent estimates of endosymbiont population size. Front Microbiol 13:816608.
74. Gu X et al. 2024. The endosymbiont *Rickettsiella viridis* increases the virulence of *Diuraphis noxia* but reduces alate frequency. bioRxiv. 2024.09.15.613162.
75. Ward S et al. 2020. Biology, ecology and management of Diuraphis noxia (Hemiptera: Aphididae) in Australia. Austral Entomol 59:238–252.
76. Dorai APS et al. 2024. Novel transinfections of Rickettsiella do not affect insecticide tolerance in Myzus persicae, Rhopalosiphum padi, or Diuraphis noxia (Hemiptera: Aphididae). J Econ Entomol 117:1377–1384.
77. Niepoth N, Ellers J, Henry LM. 2018. Symbiont interactions with non-native hosts limit the formation of new symbioses. BMC Evol Biol 18:27.
78. Yazdani M et al. 2018. First detection of Russian wheat aphid Diuraphis noxia Kurdjumov (Hemiptera: Aphididae) in Australia: a major threat to cereal production. Austral Entomol 57:410–417.
79. Chen DQ, Montllor CB, Purcell AH. 2000. Fitness effects of two facultative endosymbiotic bacteria on the pea aphid, Acyrthosiphon pisum, and the blue alfalfa aphid, A. kondoi. Entomol Exp Appl 95:315–323.
80. Lee SF, White VL, Weeks AR, Hoffmann AA, Endersby NM. 2012. High-throughput PCR assays to monitor Wolbachia infection in the dengue mosquito (Aedes aegypti) and Drosophila simulans. Appl Environ Microbiol 78:4740–4743.
81. Damsteegt VD, Gildow FE, Hewings AD, Carroll TW. 1992. A clone of the Russian wheat aphid (Diuraphis noxia) as a vector of barley yellow dwarf, barley stripe mosaic, and brome mosaic viruses. Plant Dis 76:1155–1160.
82. Aldesuquy H, Baka Z, Mickky B. 2014. Kinetin and spermine mediated induction of salt tolerance in wheat plants: leaf area, photosynthesis and chloroplast ultrastructure of flag leaf at ear emergence. Egypt J Basic Appl Sci 1:77–87.

**Table 1. Summary of effects of *Rickettsiella* and *Regiella* transinfections in *Diuraphis noxia***

| **Trait** | **Details** | ***Rickettsiella*** | ***Regiella*** |
| --- | --- | --- | --- |
| **Plant damage** | **Plant leaf number and area** | ↓ | ↑ |
|  | **Plant damage score** | ↑ | ↓ |
|  | **Chlorotic streaking** | ↑ | ↓ |
| **Plant defense** | **Plant JA levels** | No effect | No effect |
|  | **Plant SA levels** | No effect | No effect |
| **Aphid population** | **Aphid population growth** | ↑ | ↓ |
|  | **Alate production** | ↓ | No effect |
| **Endosymbiont spread** | **Frequency of endosymbiont** | Increase at 25°C | Increase at 19°C |
| **Aphid individual fitness** | **Development time, fecundity, longevity, body size, body color** | Faster development at 19°C and no other fitness effects | No effects detected on individual fitness traits |

↑ represents increase and ↓ represents decrease.

All traits are presented as comparisons relative to wildtype aphids.
